# Cystinosin/Ers1 functions in redox homeostasis in the early secretory pathway

**DOI:** 10.64898/2026.03.01.708828

**Authors:** Julia Zhu, Sumanth Mosale Seetharam, Jade Bowerman, Solomon Baik, Katie Munechika, Nathan Wu, Aleksandra Skirycz, Haiyuan Yu, Miguel Piñeros, Richa Sardana

## Abstract

Cystinosis is an autosomal recessive inherited disorder caused by mutations in the *CTNS* gene, which encodes the highly conserved transmembrane protein cystinosin, a proton/cystine co-transporter at the lysosome membrane. However, reduction of cystine load in the lysosomes is insufficient to treat key disease symptoms, indicating that cystinosin performs additional disease-relevant functions. Here, we report that Ers1, the yeast homolog of cystinosin, localizes to and functions in the early secretory pathway. We provide evidence that Ers1 does not transport cystine. Ers1 genetically interacts with early secretory pathway recycling adaptors and redox-active Fe-S cluster-binding proteins. Notably, cystinosin-LKG, the extra-lysosomal localized splicing isoform of cystinosin, can functionally replace Ers1 in yeast. Collectively, our work uncovers a conserved role of cystinosin/Ers1 in the early secretory pathway, offering new molecular insights for understanding cystinosis pathology.

## Introduction

Cystinosis is a rare, monogenic, autosomal recessive disorder caused by mutations in the *CTNS* gene encoding cystinosin. 95% of cystinosis patients present with the most severe form, referred to as infantile nephropathic cystinosis, with manifestations of photophobia, growth retardation, neuromuscular, endocrine, and kidney dysfunction progressing to end-stage kidney failure within the first decade of life (Cherqui and Courtoy, 2017; David et al., 2019; Elmonem et al., 2016). Patients with cystinosis were known to build up high levels of cystine in their lysosomes long before it was discovered that mutations in the *CTNS* gene cause the disease. (Gahl et al., 1982; Schneider et al., 1967; Town et al., 1998). Cystinosin mediates proton-dependent transport of cystine from the lysosome lumen to the cytosol (Kalatzis et al., 2001a; Ruivo et al., 2012). As a result, reduction of lysosomal cystine levels using an aminothiol, cysteamine, has been the mainstay for cystinosis therapy (Besouw et al., 2013; Jeźégou et al., 2012; Liu et al., 2012; Pisoni et al., 1985).

Cysteamine acts by reducing lysosomal cystine levels and as an antioxidant by directly scavenging free radicals (Carneiro and Jones, 2026; Jamalpoor et al., 2021; Wilmer et al., 2011). While treatment with cysteamine slows disease progression, it cannot halt or reverse kidney damage (Cherqui, 2012; Cherqui and Courtoy, 2017). Importantly, in addition to cystine transport, multiple cellular processes are significantly dysregulated by the lack of cystinosin, including autophagy (Festa et al., 2018; Luciani et al., 2018), chaperone mediated autophagy (Napolitano et al., 2015), mTOR signaling and TFEB function (Andrzejewska et al., 2016; Berquez et al., 2023; Luciani and Devuyst, 2024; Rega et al., 2016), endoplasmic reticulum homeostasis (Gaide Chevronnay et al., 2015; Johnson et al., 2013; Wei et al., 2008), redox homeostasis and glutathione metabolism (Chol et al., 2004; Mannucci et al., 2006; Sumayao et al., 2016; Wilmer et al., 2005), mitochondrial function (Levtchenko et al., 2006), inflammation (Lobry et al., 2019), and increased apoptosis (Sansanwal et al., 2010). Many of these defects are not corrected by cysteamine treatment (Cherqui, 2012; Cherqui and Courtoy, 2017). Evidence for a cystine-independent role of cystinosin is further supported by the rescue of trafficking of the chaperone-mediated autophagy receptor LAMP2A by the transport-defective CTNS-K280R mutant (Zhang et al., 2017). Clearly, the molecular causes underlying many of the defects in cystinosis require further investigation.

Cystinosin is a highly conserved heptahelical membrane protein and belongs to the PQ-loop family, members of which function as trafficking receptors, amino acid transporters, lipid flippases, or sugar transporters (Saudek, 2012). PQ-loop family proteins function in diverse essential pathways, and their deficiency is frequently associated with human disease or embryonic lethality (Anand et al., 2001; Cela et al., 2022; Jain and Zoncu, 2021; Schenk et al., 2001; Wang et al., 2022). The best characterized member of this family, KDELR, a receptor for COPI-adaptor dependent retrieval of lumenal ER resident proteins from Golgi back to the ER, has been proposed to have evolved from a transporter to a receptor (Bräuer et al., 2019; Capitani &Sallese, 2009; Kriel et al., 2011; Newstead & Barr, 2020). Some members of the PQ-loop family have been proposed to play dual roles as solute transporters as well as cargo trafficking receptors (Saudek, 2012; Stokes et al., 2005; Talaia et al., 2021). Cystinosin, too, has been proposed to play a potential role as a cargo receptor or regulator of cargo trafficking (Andrzejewska et al., 2016; Khare et al., 2025; Saudek, 2017, 2012; Zhang et al., 2017).

Cystinosin is expressed as two isoforms due to alternative splicing in exon 12- the canonical isoform, and the longer isoform referred to as CTNS-LKG that lacks the C-terminal tyrosine motif required for lysosomal targeting of the canonical isoform (Bellomo et al., 2016; Taranta et al., 2012, 2008). While the CTNS-LKG isoform is less-studied than the canonical isoform, overexpression studies have reported its localization to the lysosomes, plasma membrane, Golgi apparatus, endoplasmic reticulum (ER), and cytosolic vesicles (Bellomo et al., 2016; Taranta et al., 2016, 2008). The functional relevance of the extra-lysosomal localization of CTNS-LKG is unclear. CTNS-LKG is highly expressed in tissues associated with the clinical manifestations of cystinosis, including renal proximal tubular cells (renal Fanconi syndrome), pancreatic islets of Langerhans (diabetes mellitus), Leydig cells of the testis (male infertility), and melanocytes and keratinocytes (hypopigmentation), suggesting a role for this isoform in disease pathophysiology (Taranta et al., 2012).

Ers1 (<u>ER</u>D <u>s</u>uppressor <u>1</u>), the yeast homolog of cystinosin, was identified as a multicopy suppressor of loss of Erd1, an early Golgi protein involved in ER and Golgi resident protein maintenance (Hardwick and Pelham, 1990; Sardana et al., 2021). While this genetic interaction pointed to a possible role for Ers1 at the ER or Golgi, this interpretation was inconsistent with the established lysosomal role of cystinosin. In this work, using fluorescence microscopy, proteomics, and genetic assays, we provide evidence that Ers1 indeed localizes and functions in the early secretory pathway. Ers1 function is linked to redox homeostasis as demonstrated by its genetic interaction with Fe-S cluster-binding glutaredoxin proteins Grx6 and Grx7 in the ER/Golgi. Using targeted metabolomics and two-voltage clamp electrophysiology, we find that Ers1 does not transport cystine. Finally, Ers1 function can be complemented by the CTNS-LKG isoform, suggesting functional conservation, and providing clues to extra-lysosomal functions of CTNS-LKG.

## Results

### Ers1, the yeast homolog of cystinosin, localizes to the early Golgi

Ers1 is evolutionarily conserved, with the highest conservation in the functionally important PQ loops (**Figure 1A**). Ers1 closely aligns with the core transmembrane regions of human cystinosin, but lacks the large luminal N-terminal domain and the C-terminal tyrosine motif observed in cystinosin (**Figure 1B**). Since the C-terminal tyrosine motif is required for lysosomal targeting of cystinosin, it prompted us to carefully examine the localization of Ers1 in yeast. Endogenously tagged Ers1-GFP exhibited a punctate localization pattern and did not co-localize with the vacuolar (yeast lysosome) marker Vph1 (**Figure 1C**). A previous study using the methylotrophic yeast *Pichia pastoris*, reported an interaction between Ers1S and an endosomal protein, Nhx1, in a bimolecular fluorescence complementation assay (Khare et al., 2025). However, we observed very little overlap between endogenously tagged Ers1-GFP and Nhx1-mCherry, suggesting that Ers1 does not predominantly localize to endosomes at steady state (**Figure 1C**). We next examined the co-localization of Ers1 with early (Mnn9), medial (Gea2), and late Golgi (Sec7) markers. Interestingly, Ers1 largely co-localized with the early-Golgi marker, Mnn9, and to some extent with the medial-Golgi marker, Gea2, suggesting it functions in early Golgi compartments (**Figure 1D, 1E**). Endogenous Ers1 expression is relatively low, so we also compared the localization of Ers1-GFP expressed from CPY and GPD promoters. In all conditions, the punctate signal co-localized with Mnn9 at the early Golgi with diffuse ER staining that co-localized with DsRed-HDEL. Expression of Ers1 from GPD promoter also resulted in degradation of the fusion protein as seen by accumulation of GFP fluorescence in the vacuole (**Figure S1A**). To assess whether the C-terminal tag interfered with protein function, we tested the ability of multicopy Ers1-GFP or untagged Ers1 to suppress the sensitivity of the *erd1*Δ mutant to the aminoglycoside hygromycin (Hardwick and Pelham, 1990; Sardana et al., 2021). Both untagged Ers1 and Ers1-GFP expressed from a multicopy plasmid were able to rescue the *erd1*Δ mutant, suggesting that the C-terminal tagged protein is functional (**Figure S1B**). Furthermore, the tag position did not influence localization, as N-terminally tagged GFP-Ers1 also exhibited a punctate pattern at steady state that co-localized with the early-Golgi marker, Mnn9 (**Figure S1C**). Take together, our findings suggest that Ers1 localizes to the early Golgi at steady state.

**Figure 1.**
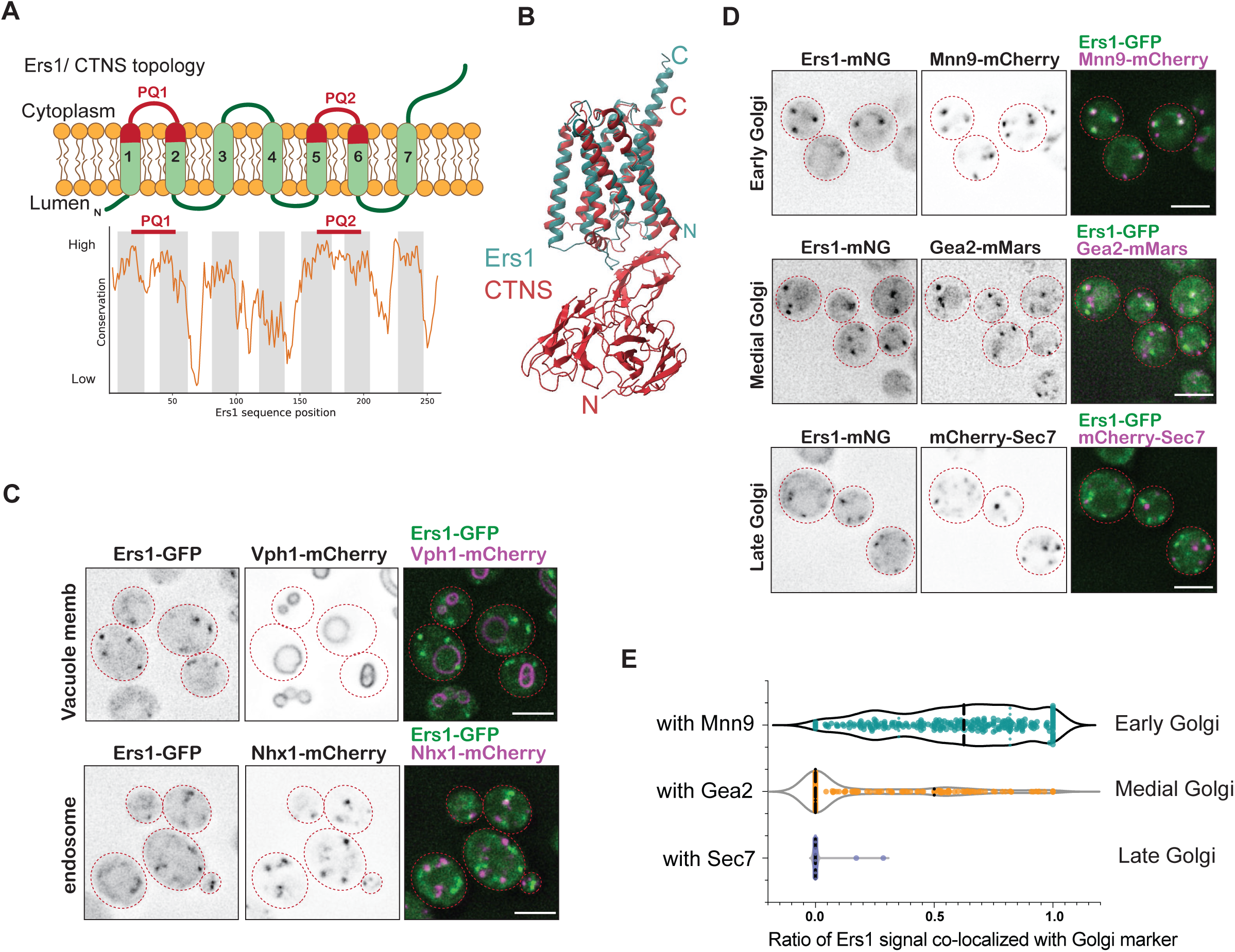
Ers1, the yeast homolog of CTNS, localizes to the early Golgi. (A) Predicted membrane topology of Ers1 and level of protein sequence conservation. Gray regions indicate predicted transmembrane domains. (B) Alignment of the structure of CTNS (red) (PDB: 8dkm) (Guo et al., 2022) with the structure of Ers1 (green) predicted by Alphafold3. (C) Live-cell fluorescence imaging of Ers1-GFP with vacuole membrane marker (Vph1-mCherry) and endosome marker (Nhx1-mCherry). Red dashed lines indicate the cell boundaries based on DIC images. Scale bar 5 µm. (D) Live-cell fluorescence imaging of Ers1-GFP with early (Mnn9-mCherry), medial (Gea2-mMars), and late (mCherry-Sec7) Golgi markers. (E) Violin plots for the ratio of co-localized Ers1-mNeonGreen fluorescence with early (Mnn9-mCherry), medial (Gea2-mMars), and late (Sec7-6xDsRed) Golgi markers (n = 300 puncta for each condition).

### Recycling by COPI vesicle coat and cargo adaptor Vps74 maintains Ers1 at the Golgi

To investigate the mechanisms underlying Ers1 localization, we first examined Ers1-GFP localization in COPII loss-of-function temperature-sensitive (ts) mutant. The conserved cytoplasmic COPII coat complex mediates cargo sorting from ER to the Golgi (Downes and Zanetti, 2025; Lord et al., 2013). Under permissive conditions (26 °C), Ers1-GFP exhibited weak ER and punctate localization in the *sec23-1* COPII mutant background. Upon shift to restrictive temperature (38 °C) for 30 min, Ers1 was completely retained in the ER in the *sec23-1* mutant, supporting a requirement for COPII-dependent vesicle trafficking in its ER-to-Golgi transport (**Figure 2 A, S2A**).

**Figure 2.**
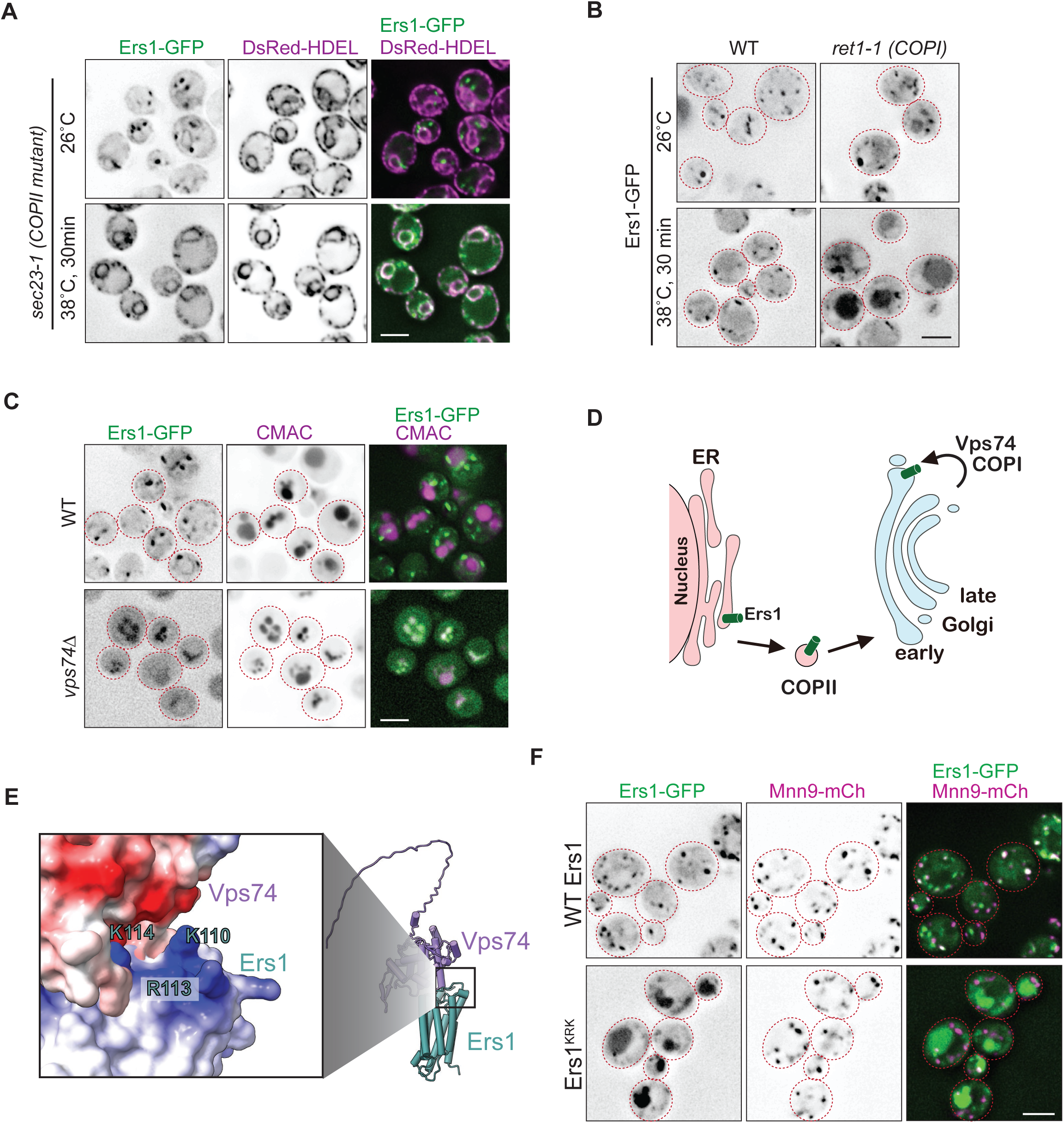
Recycling by COPI vesicle coat and cargo adaptor Vps74 maintains Ers1 at the Golgi. (A) Localization of Ers1-GFP in the COPII ts mutant *sec23-1*. Cells were grown to mid-log phase at 26°C and then incubated at 38°C for 30 min before imaging. Scale bar 5 µm. (B) Localization of Ers1-GFP in the COPI ts mutant *ret1-1*. Cells were grown to mid-log phase at 26°C and then incubated at 38°C for 30 min before imaging. (C) Localization of Ers1-GFP in wild-type and *vps74*Δ. Cells were grown in a synthetic medium and vacuoles were stained with CMAC. (D) Schematic showing Ers1 is trafficked from the ER to the Golgi by COPII and recycled at the Golgi by COPI and Vps74. (E) Structure of an Ers1 and Vps74 complex predicted by Alphafold2, showing interaction of a basic (blue) cytosolic loop in Ers1 (K110A, R113A, K114A) with the acidic (red) surface of Vps74. (F) Localization of wild-type and mutant (K110A, R113A, K114A) Ers1-GFP in wild-type cells with an early Golgi marker (Mnn9-mCherry).

The distribution of Ers1 at the ER and Golgi at steady state suggests that Ers1 is maintained in the early secretory pathway by recycling mechanisms. To probe whether Ers1 is recycled back from the endosome or later Golgi cisternae, we examined the localization of Ers1-GFP in loss of function mutants of PI 3-kinase Vps34 (*vps34-ts*), AP-1 adaptor (*apl2*Δ), and COP1 (*ret1-1* ts). While the localization of Ers1 was unaffected in Vps34 or AP-1 mutants, it was mislocalized to the vacuole lumen in the *ret-1* COPI mutant at non-permissive temperature, suggesting that COPI function is required to maintain Ers1 at the Golgi (**Figure 2B, S2B**). Many early Golgi resident transmembrane proteins depend on the cytosolic cargo adaptor of COPI, Vps74, for recycling (Sardana et al., 2021; Schmitz et al., 2008; Tu et al., 2008). Indeed, deletion of Vps74 also caused Ers1-GFP to mislocalize to the vacuole lumen (**Figure 2C**). These results suggest that Ers1 is maintained in the early-secretory pathway by COPII dependent anterograde traffic and Vps74-COPI dependent retrograde traffic (**Figure 2D**). COPI and Vps74 are known to bind basic residues in the cytosolic region of resident Golgi proteins for their recycling (Letourneur et al., 1994; Schmitz et al., 2008; Taylor et al., 2025; Tu et al., 2008). We identified and mutated several basic patches in the cytosolic regions of Ers1 and found that a positively charged region in the cytosolic loop between transmembrane domains 3 and 4 of Ers1, comprised of residues K110, R113, and K114 is required for the correct localization of Ers1 (**Figure S2C**). Alphafold modeling (Mirdita et al., 2022) of Vps74 and Ers1 predicts this positively charged region to form an interaction surface with the negatively charged region of Vps74 previously demonstrated to interact with cargos (Taylor et al., 2025) (**Figure 2E**). Co-localization analysis indicated *Ers1^KRK^* mutant was not maintained at the early Golgi (**Figure 2F**) and was instead delivered to the vacuole lumen for degradation (**Figure S2D**). Collectively, these findings establish the mechanistic basis for the maintenance of Ers1 at the early Golgi by COPII dependent anterograde and Vps74-COPI dependent retrograde trafficking.

### Ers1 physically interacts with ER and Golgi proteins and is a EGAD target

Our localization studies suggested a role for Ers1 in the early secretory pathway. To explore this in an unbiased manner, we sought to identify proteins in close proximity to Ers1. We utilized Ers1-GFP expressed from a GPD promoter for protein immunoprecipitation followed by mass spectrometry. Gene ontology analysis on the significant hits revealed a strong enrichment of ER and Golgi proteins immunoprecipitated with Ers1 (**Figure 3A, 3B**). Notably, all subunits of the Dsc E3 ligase complex (Tul1, Dsc2, Dsc3, Ubx3, Cdc48) were significantly enriched in the Ers1 interactome dataset (**Figure 3A**). Dsc complex and Tul1 E3 ligase complex have been reported to function in membrane protein ubiquitination and quality control at the Golgi to facilitate Endosome and Golgi-associated Degradation (EGAD) (Schmidt et al., 2019; Weyer et al., 2024). To validate the enrichment of the Dsc complex with Ers1, we repeated the Ers1-GFP IP followed by western blotting. Indeed, all the components of the Dsc complex were enriched in the IP (**Figure 3C**). To assess if Ers1 might play a role in EGAD, we monitored the stability of an EGAD substrate, Orm2, in wild-type yeast and in mutants lacking Ers1, Tul1, or both. While loss of Tul1 E3 ligase resulted in the stabilization of FLAG-Orm2, we did not observe an effect due to loss of Ers1, suggesting Ers1 is likely not required for EGAD (**Figure 3D**). We next asked if Ers1 itself might be an EGAD target. We surmised that saturation of recycling machinery by overexpression of Ers1 might result in the overflow of Ers1 to late Golgi compartments. This pool that escapes the medial Golgi could be marked by the Dsc E3 ubiquitin ligase complex for degradation. Indeed, deletion of Tul1, E3 ubiquitin ligase of the Dsc complex, resulted in stabilization of Ers1 (**Figure 3E**). Additionally, overexpression of Tul1 resulted in downregulation of Ers1 and sorting of Ers1 to the vacuole lumen for degradation (**Figure 3F, 3G**). Overall, these results suggest that the levels and localization are monitored by the Dsc E3 ligase mediated Golgi protein quality control and identify Ers1 as a novel EGAD target.

**Figure 3.**
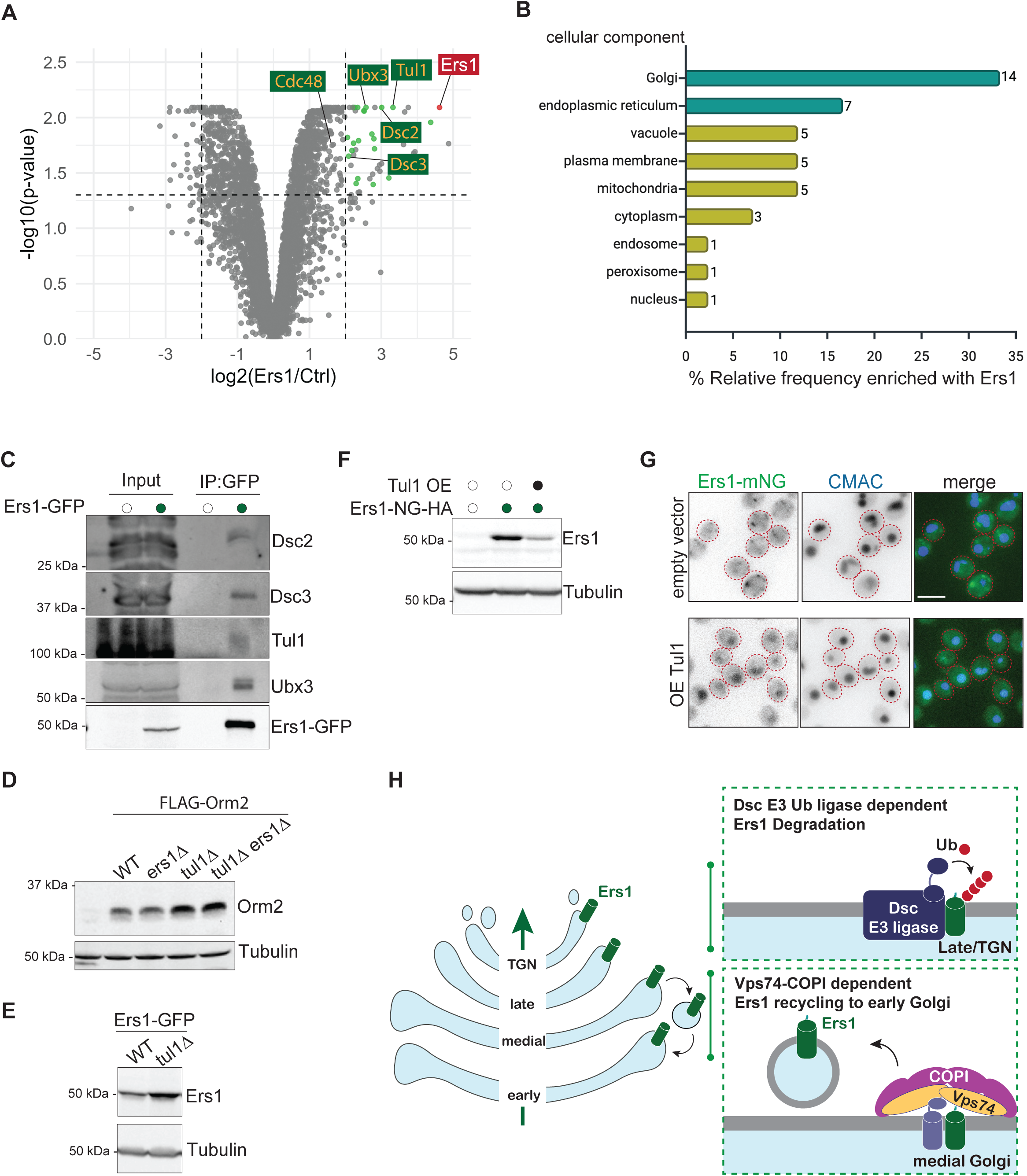
Ers1 physically interacts with ER and Golgi proteins and is a EGAD target. (A) Volcano plot of proteins identified in IP-MS of Ers1-GFP. Proteins with a t-test p value <0.05 and more than 2-fold enrichment compared to the control IP (log2FC>2) were considered to be significant. Resident ER and Golgi proteins that pass significance thresholds (dotted lines) are colored in green. The hits corresponding to Ers1 (bait) is highlighted in Red and components of Dsc E3 ligase complex in orange. (B) Relative frequency of the proteins enriched with Ers1 (log2FC>2, p value <0.05) annotated based on cellular compartment. (C) Immunoprecipitation of Dsc E3 ubiquitin ligase complex components with Ers1-GFP expressing or untagged yeast cells. (D) Western blot analysis of whole cell lysates from wild-type, *ers1Δ*, *tul1Δ*, and *ers1Δtul1Δ* cells to examine stability of EGAD target, FLAG-Orm2. (E) Western blot analysis of whole cell lysates from wild-type and *tul1*Δ cells expressing Ers1-GFP. (F) Western blot analysis of whole cell lysates from cells expressing either endogenously tagged Ers1-NG-HA or untagged Tul1 overexpressed from the Tet-off promoter. (G) Localization of Ers1-NG-HA in cells expressing an empty vector or a plasmid encoding for untagged Tul1 expressed under the Tet-off promoter. Cells were grown in a synthetic medium and vacuoles were stained with CMAC. (H) Schematic showing regulation of Ers1 localization and levels at the Golgi. Recycling by Vps74/COPI facilitates localization of Ers1 at the early-Golgi at steady state. Dsc E3 ligase complex facilitates quality control at the late-Golgi to mediate degradation of Ers1 that bypasses recycling at the early Golgi cisternae.

### Ers1 influences thiol-redox homeostasis in the early secretory pathway

A previous analysis of *ers1*Δ mutant reported transcriptional downregulation of numerous genes involved in iron and redox homeostasis (Simpkins et al., 2016) (**Figure 4A**). Notably, many of these genes play roles in glutathione homeostasis, including the GSH antiporter Gex1, glutaredoxin Grx8, and amino acid transporters of glutamine (Agp3) and cysteine (Yct1), both of which are precursors of the GSH synthesis pathway (Simpkins et al., 2016). Our observations support an early Golgi localization and function for Ers1, prompting us to examine its link to redox processes in the early secretory pathway. Although redox homeostasis is critical for ER protein folding and function (Ellgaard et al., 2018; Sevier and Kaiser, 2008), not much is known about its role or how it is maintained in the Golgi apparatus (Kellokumpu, 2019).

**Figure 4.**
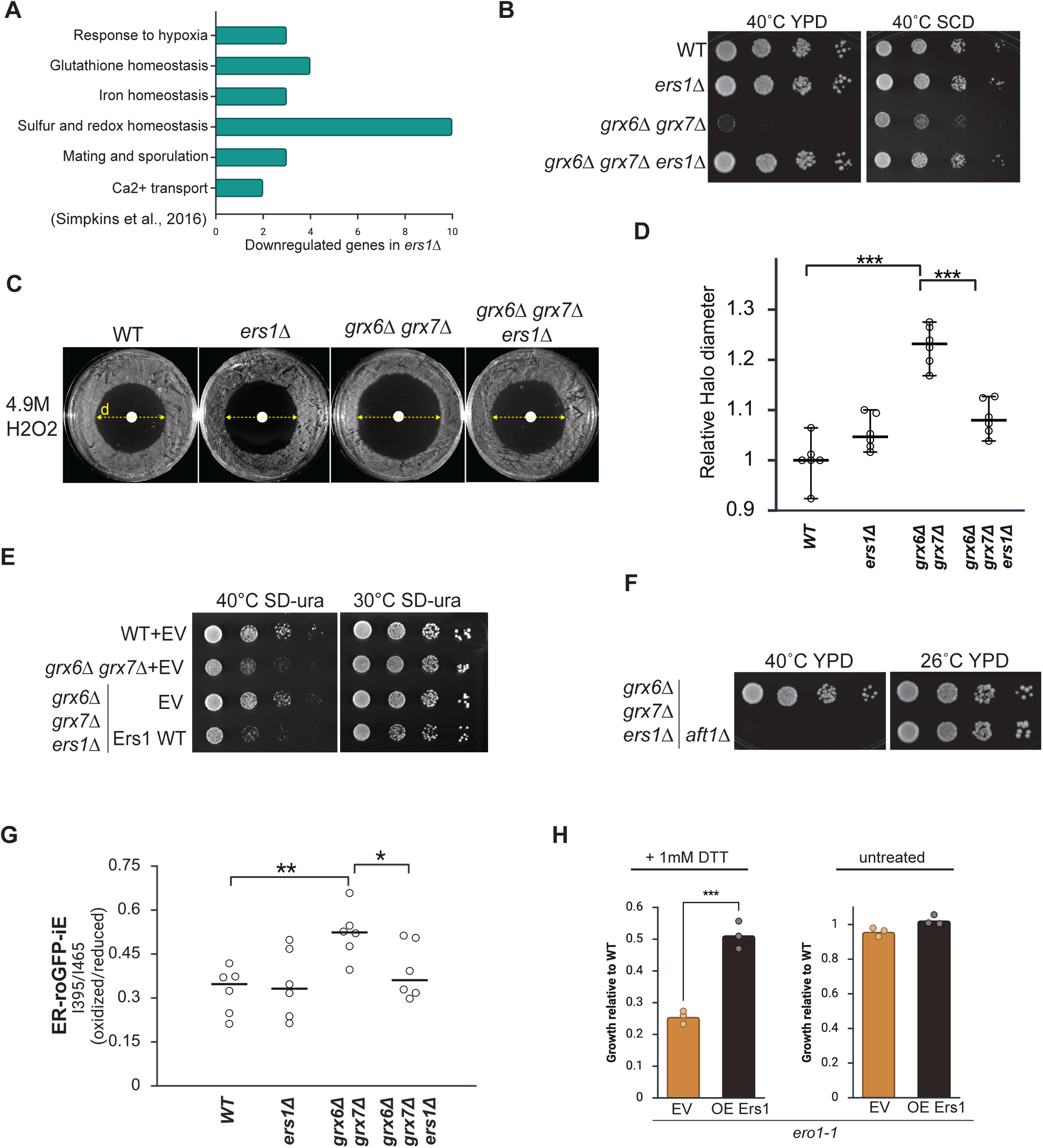
Ers1 influences ER and Golgi thiol-redox homeostasis. (A) Enriched categories of gene transcriptionally downregulated in the *ers1*Δ mutant reported in Simpkins et al., 2016. (B) Growth under temperature stress of wild type, *ers1*Δ, *grx6Δgrx7Δ*, and g*rx6Δgrx7Δers1Δ* cells. Cells were spotted as seriated dilutions on YPD or SCD plates and incubated at 40°C for 3 days. (C) Disc diffusion assay for wild-type, *ers1Δ*, *grx6Δgrx7Δ*, and *grx6Δgrx7Δers1Δ* sensitivity to hydrogen peroxide. Cultures were plated on SCD plates and filter discs soaked with 10 µl of 4.9 M H2O2 were placed at the center of each plate before incubation at 30°C. (D) Quantification of diameters of growth inhibition halos of wild-type, *ers1Δ, grx6Δgrx7Δ*, and *grx6Δgrx7Δers1Δ* cells to peroxide. Bar graphs represent averages from three independent experiments, shown as individual dots. ***, p<0.001 (two-tailed Student’s t-test). (E) Growth under temperature stress of wild-type, *grx6Δgrx7Δ*, and *grx6Δgrx7Δers1Δ* cells transformed with either an empty vector or a plasmid expressing wild-type Ers1. Cells were spotted as seriated dilutions on SD-ura plates and incubated at 30°C or 40°C for 3 days. (F) Growth under temperature stress of *grx6Δgrx7Δers1Δ* and *grx6Δgrx7Δers1Δaft1Δ* cells. Cells were spotted as seriated dilutions on YPD plates and incubated at 40°C or 26°C for 3 days. (G) Measurements of ER redox state using ER-roGFP-iE reporter in wild-type, *ers1Δ*, *grx6Δgrx7Δ*, and *grx6Δgrx7Δers1Δ* cells. Bar graphs represent averages from five independent experiments, shown as individual dots. *, p<0.05, **, p<0.01 (two-tailed Student’s t-test). (H) Growth of wild type and *ero1-1* cells transformed with an empty vector or plasmid overexpressing Ers1 untreated or upon treatment with 1 mM DTT. Cells were grown in synthetic media for 24 hours and growth of *ero1-1* mutant was normalized with wild type for the respective treatment. Bar graphs represent averages from three independent experiments, shown as individual dots. ***, P<0.001 (two-tailed Student’s t-test).

Two putative Fe-S cluster binding glutathione-dependent oxidoreductases or glutaredoxins, Grx6 and Grx7, have been previously reported to function in the ER and early-Golgi (Izquierdo et al., 2008; Luo et al., 2010; Mesecke et al., 2008). Loss of Grx6 and Grx7 mutants display an increased sensitivity toward oxidizing agents such as peroxide and temperature stress, highlighting their role in oxidative stress resistance (Mesecke et al., 2008). To investigate if Ers1 plays a role in redox homeostasis at the Golgi, we examined its genetic interaction with Grx6 and Grx7. As reported previously, yeast lacking Grx6 and Grx7 were highly sensitive to temperature stress and hydrogen peroxide (**Figure 4B, S4A, 4C, 4D**). Strikingly, deletion of Ers1 robustly suppressed the temperature and hydrogen peroxide sensitivity of the *grx6Δ grx7Δ* double mutant (**Figure 4B, 4C, 4D**). This suppression was dependent on loss of Ers1 function as expression of Ers1 on a plasmid restored the temperature sensitivity (**Figure 4E**). Loss of Grx6 and Grx7 did not alter the punctate localization of Ers1 suggesting the phenotype is not due to Ers1 mislocalization (**Figure S4B**). Disruptions in thiol redox balance have previously been reported to activate the Aft1 transcription factor, leading to remodeling of iron metabolism (Outten and Albetel, 2013). Oxidative stress, glutathione levels, as well as activity of cytosolic glutaredoxins Grx3/4, are linked to the transcriptional activation of Aft1-dependent responses (Kumar et al., 2011; Pujol-Carrion et al., 2006). Suppression of temperature sensitivity of *grx6Δ grx7Δ* upon deletion of Ers1 was dependent on Aft1 function, suggesting the response is dependent on oxidative stress induced changes (**Figure 4F**).

To understand the influence of Ers1 on the redox environment of the Golgi, we targeted a ratiometric redox sensor, roGFPiE (Lohman and Remington, 2008), to the early Golgi in yeast by fusing the sensor to the luminal domain of early Golgi glycosyltransferases, Och1 and Hoc1. However, the signal-to-background ratio in Golgi-roGFPiE expressing yeast cells was very low, preventing reliable signal detection. Given that Ers1, Grx6, and Grx7 also show ER localization (Mesecke et al., 2008), we decided to use a previously characterized ER-localized roGFP-iE probe (Wang and Sevier, 2016) to examine the contribution of Ers1 to the ER redox state. Loss of Grx6 and Grx7 resulted in a significant increase in oxidized ER-roGFP-iE, and deletion of Ers1 suppressed this oxidizing effect (**Figure 4G**). These results suggest that Ers1 activity may contribute to increased oxidation in the ER/Golgi that is detrimental in the absence of Grx6 and Grx7. Ero1 is an essential ER protein disulfide oxidase that enables oxidative protein folding in the ER (Frand and Kaiser, 1998; Pollard et al., 1998). Mutations that reduce the activity of Ero1 reduce the oxidizing environment and are hypersensitive to reducing agents such as DTT (Frand and Kaiser, 1998; Pollard et al., 1998). Previous work has reported that the DTT sensitivity of the *ero1-1* mutant can suppressed by the deletion of Grx6 (Puigpinós et al., 2015). Consistent with a more oxidizing role, overexpression of Ers1 in an *ero1-1* mutant partially suppressed DTT sensitivity (**Figure 4H**). Taken together, these results establish a role of Ers1 in redox regulation in the ER/Golgi.

### Ers1 does not transport cystine at the Golgi

Cystinosin is well characterized for its cystine transport function (Kalatzis et al., 2001b; Ruivo et al., 2012). Ers1-dependent cystine transport using an ‘inside-out’ vacuole model was reported but *ers1Δ* cells failed to accumulate intracellular cystine (Simpkins et al., 2016). Since Ers1 localizes at the early-Golgi, we asked whether Ers1 transports cystine across the Golgi membrane. We hypothesized that if Ers1 transports cystine at the Golgi, then *ers1*Δ mutants would accumulate cystine in the Golgi lumen, potentially resulting in its secretion into the extracellular medium via constitutive exocytosis (**Figure 5A**). We tagged an early Golgi protein, Erd1, with GFP and optimized a workflow for rapid immunoisolation of intact early-Golgi (**Figure S5A, S5B**). Next, we employed targeted metabolomics to measure cystine and cysteine levels in total cells, immunoisolated Golgi, and secreted media from wild type or *ers1*Δ mutant. Neither cystine nor cysteine was detected in the isolated Golgi from either the wild-type or *ers1Δ* cells, despite detection of chemical standards by mass spectrometry. Since the cytosol is more reducing, we readily detected cysteine in total cell lysates and cystine in the culture supernatant. However, there was no difference in the quantified levels either between wild type and *ers1Δ* mutant (**Figure 5B, 5C**). Thus, in the absence of Ers1, yeast cells neither accumulated cystine in the Golgi, total cell, or secreted media. Since glutathione is an important redox buffer in the early secretory pathway (Birk et al., 2013), we tested whether deletion of Ers1 impacts the levels of reduced (GSH) and oxidized (GSSG) glutathione. As has been reported for the ER, the GSH∶GSSG ratio in the Golgi was lower than the cytosol, suggesting a more oxidizing Golgi environment (Hwang et al., 1992). However, wild-type and *ers1Δ* mutant did not differ in GSH or GSSG levels in isolated Golgi, total cell, or the culture supernatant (**Figure 5D, 5E**).

**Figure 5.**
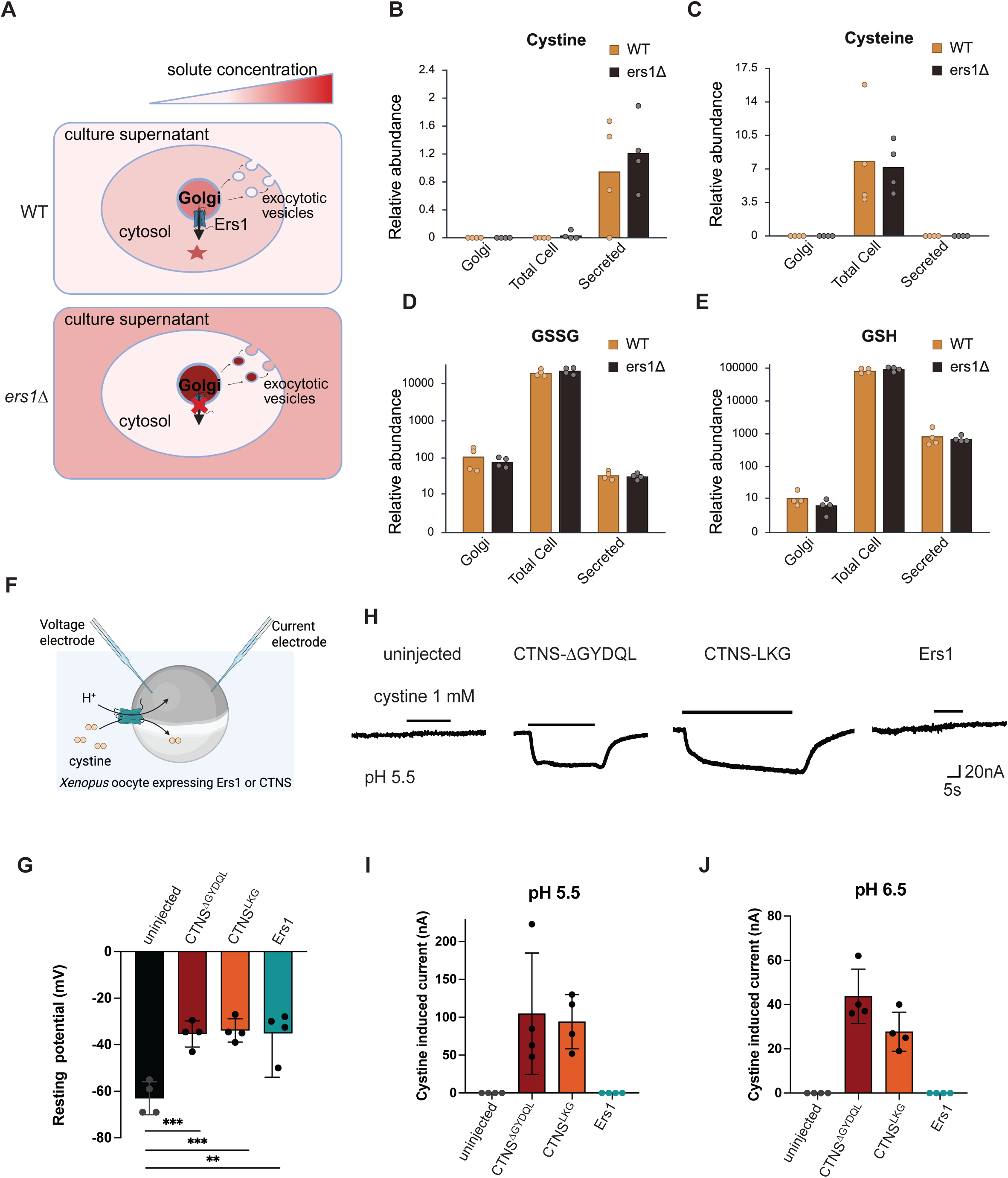
Ers1 does not transport cystine. (A) Schematic showing the prediction for cystine accumulation in the Golgi lumen or in the extracellular media in *ers1*Δ mutant for metabolomics. (B) and (C) Quantification of cystine (B) and cysteine (C) in immunoisolated Golgi, total cell lysate, and culture supernatant of wild-type and *ers1*Δ cells. Bar graphs represent averages from four biological replicates, shown as individual dots. (D) and (E) Quantification of oxidized (GSSG) (D) and reduced glutathione (GSH) (E) in immunoisolated Golgi, total cell lysate, and culture supernatant of wild-type and ers1Δ cells. Bar graphs represent averages from four biological replicates, shown as individual dots. (F) Schematic of Two-electrode Voltage Clamp (TEVC) setup in Xenopus oocytes to measure transport of cystine. (G) Resting membrane potentials of uninjected oocytes and those expressing CTNS-ΔGYDQL, CTNS-LKG, and Ers1 in standard bath solution at pH 6.5. Bar graphs represent averages from four biological replicates, shown as individual dots. (H) Representative currents induced by the addition of 1 mM cystine at pH 5.5 to the basal solution bathing uninjected oocytes and oocytes expressing CTNS-ΔGYDQL, CTNS-LKG, and Ers1. Currents were recorded under voltage clamp conditions at a holding potential of -80mV. (I) and (J) Quantification of cystine induced currents in uninjected oocytes and oocytes expressing CTNS-ΔGYDQL, CTNS-LKG, and Ers1 at pH 5.5 (I) and pH 6.5 (J). Bar graphs represent averages from four biological replicates, shown as individual dots.

To directly test whether Ers1 can transport cystine, we performed electrophysiological measurements in *Xenopus* oocytes expressing Ers1 or CTNS isoforms (**Figure 5F**). As a positive control for transport, we expressed cystinosin lacking the C-terminal lysosomal targeting motif (GYDQL) as it was previously reported to increase surface expression in oocytes (Ruivo et al., 2012). Fluorescently tagged Ers1, CTNS^ΔGYDQL^, and CTNS^LKG^ could be detected at the oocyte surface, indicating plasma membrane targeting (**Figure S5C**). Compared with uninjected control cells, oocytes expressing untagged Ers1, CTNS^ΔGYDQL^, and CTNS^LKG^ displayed depolarized resting membrane potentials when impaled in standard ND96 bath solution (**Figure 5G**). These observations suggested that these proteins encode functional transporters that can increase net cation influx, thereby reducing the net internal negative charge, consistent with the previously reported proton co-transport mechanism (Ruivo et al., 2012). Under two-electrode voltage clamp (TEVC), addition of cystine to the extracellular medium at pH 5.5 (lysosomal pH) or pH 6.5 (early Golgi pH) induced an inward (negative) current in cells expressing CTNS^ΔGYDQL^ and CTNS^LKG^, with the currents being larger at the more acidic pH. In contrast, no cystine induced current was observed in oocytes expressing Ers1 (**Figure 5H, S5D, 5I, 5J**). Our *in vivo* metabolomics data in yeast did not detect cystine in the Golgi lumen. However, glutathione, a cysteine-containing tripeptide and major cellular redox metabolite, is abundant in the early secretory pathway (Birk et al., 2013). We therefore also tested cysteine as well as reduced and oxidized glutathione in the TEVC assay. However, no induced currents were observed in oocytes expressing CTNS^ΔGYDQL^, CTNS^LKG^, or Ers1 for any of these solutes. Collectively, our findings from metabolomics and electrophysiology experiments suggest that Ers1 is unlikely to transport cystine across the Golgi membrane *in vivo*.

### Cytosolic tail of Ers1 is important for its Golgi function

The inability of Ers1 to transport cystine is surprising since key cystine-binding residues in cystinosin are conserved in Ers1 (Guo et al., 2022; Löbel et al., 2022) (**Figure 6A**). If Ers1 transports a solute other than cystine, the cystine-binding residues may still be important for this activity. Since loss of Ers1 function robustly suppresses the temperature sensitivity of *grx6Δ grx7Δ* mutant, we tested the effect of mutating the residues corresponding to cystine-binding in Ers1. Introduction of non-functional Ers1 is expected to improve growth fitness upon heat stress in *grx6Δ grx7Δ ers1Δ* triple mutant as compared to wild type Ers1 (**Figure 6B**). Intriguingly, all cystine-binding mutants behaved like WT Ers1 and remained functional in this assay (**Figure 6C, 6D**), suggesting that the function of Ers1 in this system is possibly independent of solute transport.

**Figure 6.**
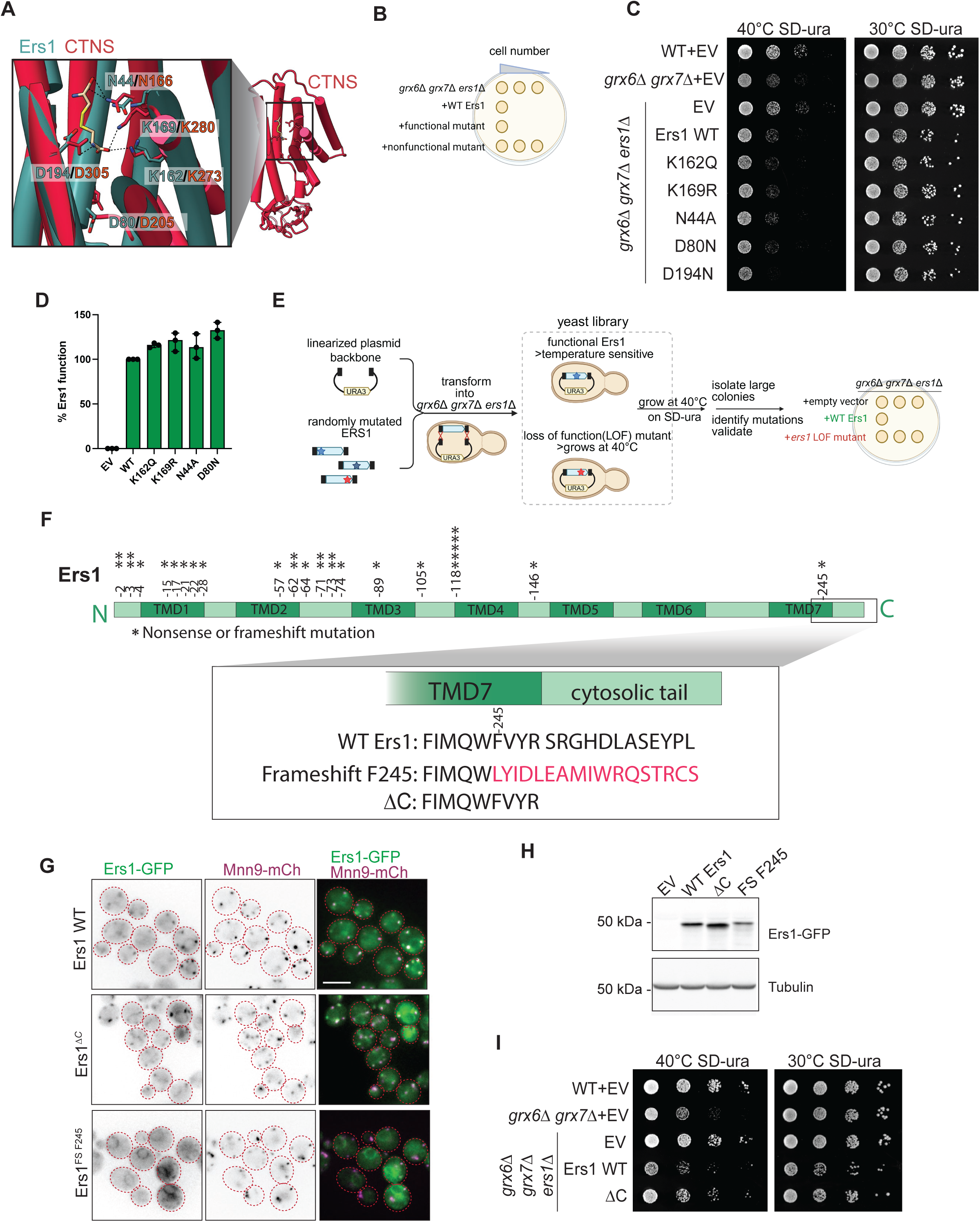
Cytosolic tail of Ers1 is important for its Golgi function. (A) Alignment of the structure of CTNS bound to cystine (red) (PDB: 8dkm) (Guo et al., 2022) with the structure of Ers1 (green) predicted by Alphafold3, with cystine-transport associated residues highlighted. (B) Schematic of expected temperature sensitivity phenotype of Ers1 complementation of *grx6Δgrx7Δers1Δ* cells. (C) Growth under temperature stress of wild-type, *grx6Δgrx7Δ*, and *grx6Δgrx7Δers1Δ* cells expressing wild-type or putative cystine-binding residues in Ers1 based on conservation with CTNS. Cells were spotted as seriated dilutions on SD-ura plates and incubated at 30°C or 40°C for 3 days. (D) Quantification of Ers1 function based on complementation of function in *grx6Δgrx7Δers1Δ* cells in (C). Bar graphs represent averages from three biological replicates, shown as individual dots. (E) Schematic for the mutagenesis screen to identify Ers1 loss of function (LOF) mutants that fail to complement the *grx6Δgrx7Δers1Δ* mutant. (F) Summary of the nonsense and frameshift mutations identified in the random mutagenesis screen. The C-terminal cytosolic tail of wildtype Ers1, and the frameshift F245 mutant identified from the screen is highlighted. The altered amino acid sequence in the mutant is highlighted in red. (G) Fluorescence microscopy analysis of GFP-tagged wild-type Ers1, ΔC, and frameshift F245 mutants in cells expressing early-Golgi marker Mnn9-mCherry. (H) Western blot analysis of GFP-tagged wild-type Ers1, ΔC, and frameshift F245 mutants. (I) Growth under temperature stress of wild-type, *grx6Δgrx7Δ*, and *grx6Δgrx7Δers1Δ* cells expressing wild-type and Ers1 ΔC mutant. Cells were spotted as seriated dilutions on SD-ura plates and incubated at 30°C or 40°C.

To identify regions of Ers1 required for its genetic interaction with Grx6 and Grx7, we generated a library of randomly mutagenized Ers1 variants using error-prone PCR and introduced them in *grx6Δ grx7Δ ers1Δ* triple mutant. Mutations that result in Ers1 loss-of-function will result in improved growth at elevated temperature (**Figure 6E**). Using this strategy, we isolated 47 loss-of-function Ers1 mutants. Most mutations were nonsense or frameshift mutations, whereas three full-length Ers1 variants harbored missense substitutions (**Figure 6F**). However, all three missense variants displayed either reduced protein stability or aberrant subcellular localization (**Figure S6A, S6B, S6C**). One frameshift mutation at F245 near the C-terminus was particularly notable. This mutation alters the sequence of the last four amino acids of the seventh transmembrane domain (TMD7) and the final 12 cytosolic amino acids (**Figure 6F**). The Ers1-F245FS variant accumulated in the ER and showed reduced steady-state stability, likely due to destabilization of the transmembrane domain and subsequent targeting by ER-associated quality control during protein folding (**Figure 6G, 6H**).

To assess the functional significance of the C-terminus, we generated a truncation mutant lacking the final 12 cytosolic residues (Ers1ΔC) (**Figure 6F**). Unlike the frameshift mutant, Ers1ΔC localized properly, co-localizing with the early Golgi marker Mnn9, and was stable at steady state (**Figure 6G, 6H**). However, Ers1ΔC was functionally defective, as indicated by improved high-temperature growth of the *grx6Δ grx7Δ ers1Δ* strain compared to wild-type Ers1 (**Figure 6I**). Because deletion of the cytosolic tail is unlikely to disrupt solute transport directly, we propose that this region mediates a protein interaction required for Ers1 function at the early Golgi.

### CTNS-LKG can complement Ers1 function at the Golgi

To test whether the role Ers1 plays in the early secretory pathway is conserved, we tested the ability of the human isoforms CTNS and CTNS-LKG to complement Ers1 function. Strikingly, CTNS-LKG, but not CTNS, was able to suppress the hygromycin sensitivity of *erd1Δ* mutant (**Figure 7A**). CTNS-LKG also complemented Ers1 function in the *grx6Δ grx7Δ* temperature sensitivity assay (**Figure 7B**). As noted earlier, the canonical isoform, CTNS, and the longer isoform, CTNS-LKG, only differ in the sequence and length of the C-terminal cytosolic tail (**Figure 7C**). CTNS-LKG isoform contains a longer cytoplasmic tail and lacks the tyrosine based lysosomal targeting motif present at its C-terminus of the canonical isoform. Unlike the canonical isoform, overexpressed CTNS-LKG has been reported to have extensive extra-lysosomal localization (Bellomo et al., 2016; Taranta et al., 2016, 2008). To assess if this is representative of endogenous expression levels of CTNS, we generated GFP tagged knock-in cell lines expressing either CTNS or CTNS-LKG isoform from the native CTNS promoter (Manna et al., 2019) (**Figure 7D**). As expected, CTNS-GFP exhibited a predominant lysosomal localization pattern (**Figure 7E**). Consistent with overexpression studies, endogenous CTNS-LKG-GFP showed a more dispersed localization pattern (Taranta et al., 2016). Compared with CTNS, CTNS-LKG showed reduced lysosomal and endosomal co-localization, increased cytosolic punctate fluorescence, and enhanced localization to the Golgi and plasma membrane. (**Figure 7E, 7F, 7G**). The extra-lysosomal localization of CTNS-LKG, and the complementation of the *ers1Δ* mutant, strongly suggest a conservation of cystine transport-independent Ers1 function outside of the lysosome. Overall, our findings identify a key gap in our knowledge of cystinosin function and underscore the importance of studying its functions beyond the lysosome.

**Figure 7.**
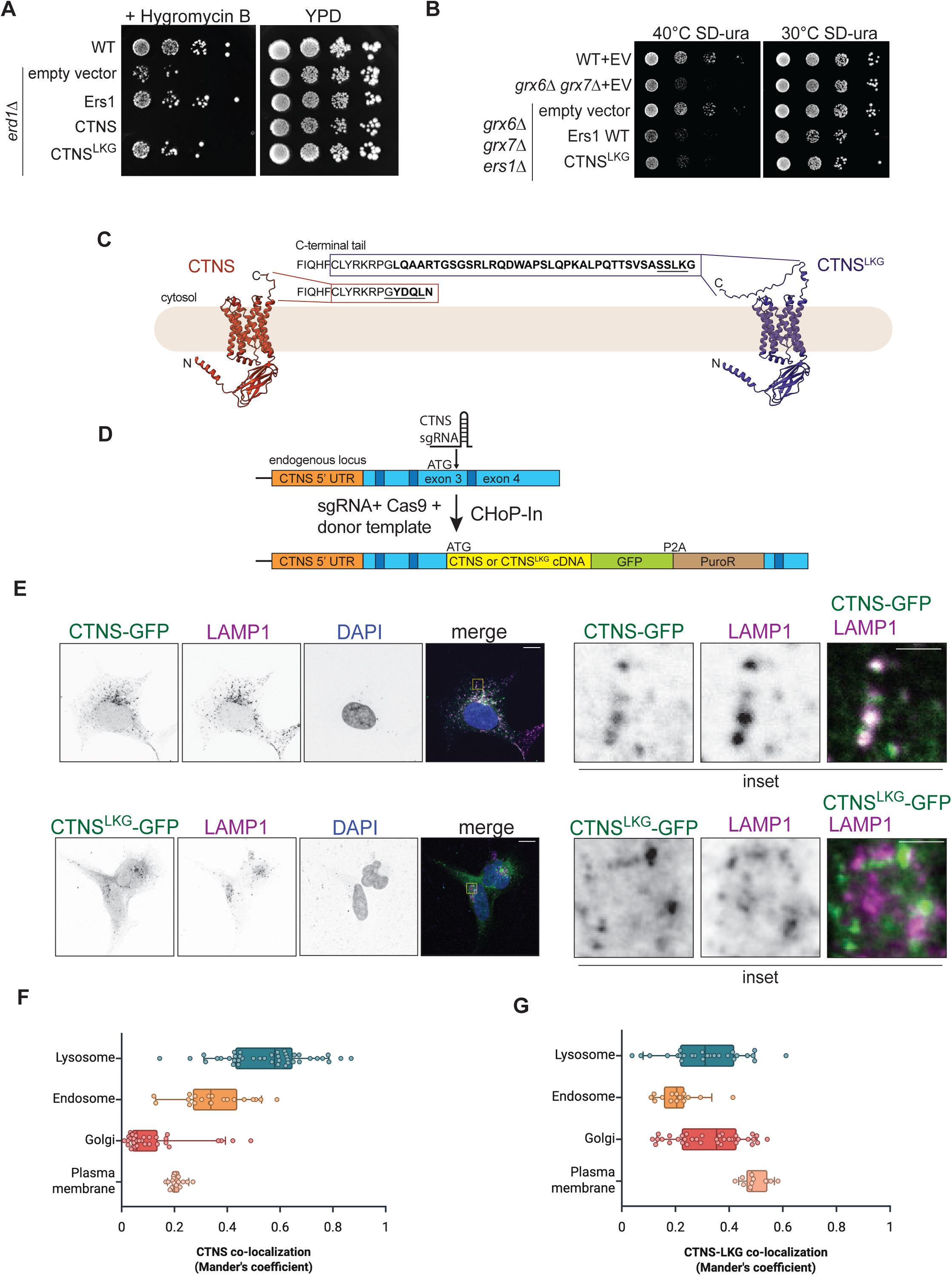
CTNS-LKG can complement Ers1 function at the Golgi. (A) Growth of cells overexpressing wild-type Ers1, human CTNS, or human CTNS-LKG on the hygromycin sensitivity of *erd1Δ* cells. Cells were spotted as serial dilutions on YPD medium with or without 25µg/ml HygromycinB for 3 days. (B) Growth under temperature stress of wild-type, *grx6Δgrx7Δ*, and *grx6Δgrx7Δers1Δ* cells expressing wild-type Ers1 or human CTNS-LKG. Cells were spotted as seriated dilutions on SD-ura plates and incubated at 30°C or 40°C for 3 days. (C) Schematic showing differences in the C-terminal amino acid sequence and length of CTNS and CTNS-LKG isoforms. (D) Schematic showing the strategy used to generate endogenously GFP tagged CTNS or CTNS-LKG expressed from the native CTNS promoter in HEK293T cells. (E) Immunofluorescence analysis of CTNS-GFP or CTNS-LKG-GFP (green) co-stained with a lysosome marker LAMP1 (red). Nuclei are counterstained with DAPI (blue). Scale bar, 10 μm. Region highlighted by yellow boxes are enlarged on the right. Scale bar, 2 µm. (F) and (G) Quantification of colocalization of CTNS-GFP (F) or CTNS-LKG-GFP (G) with organelle markers LAMP1 (lysosomes), GM130 (Golgi), EEA1 (early endosomes), and EGFR (plasma membrane) using Manders’ coefficient.

## Discussion

### Novel functional location for cystinosin in the secretory pathway

Cystinosis is a rare, debilitating inherited disorder caused by loss of cystinosin function. While cystinosin’s role as a lysosomal cystine–proton symporter is well established, accumulating evidence suggests additional functions beyond cystine transport. These include the inability of cystine depletion alone to halt disease progression, defects in cell biological pathways unrelated to cystine transport, and the extra-lysosomal localization of the CTNS-LKG isoform (Cherqui and Courtoy, 2017). Despite their relevance to disease, cystine-independent functions of cystinosin remain poorly defined. Our work challenges the assumption that the cystinosin family functions solely as lysosomal transporters and provides a new framework for understanding the non-canonical functions of cystinosin.

Our work has uncovered a novel functional role of the yeast homolog of cystinosin, Ers1, in the early secretory pathway. Despite the discovery of Ers1 as a functional interactor of Erd1, it was assumed to function at the yeast vacuole based on the known function of cystinosin as the lysosomal cystine transporter (Gao et al., 2005). It is important to note that Ers1 lacks the tyrosine motif required for the lysosomal targeting of cystinosin. Overexpressed Ers1S-GFP in *Pichia pastoris* was reported to co-localize with the lipophilic dye FM4-64, but also exhibited strong ER and non-endosomal punctate signal (Khare et al., 2025). Using endogenously tagged Ers1 expressed from its native promoter, we demonstrate that Ers1 is primarily localized to the early Golgi at steady state in budding yeast. The minor co-localization observed with the late-Golgi marker Sec7 and the endosomal marker Nhx1 likely reflects trafficking intermediates rather than steady-state localization. We demonstrate that Ers1 is maintained in the early secretory pathway by COPI and the cargo adaptor, Vps74. Indeed, overexpression of Ers1 saturated the recycling machinery, leading to increased co-localization with late-Golgi markers and accumulation in the vacuolar lumen, consistent with vacuolar degradation. Our results also indicate that the Ers1 localization at the Golgi is tightly regulated. Ers1 is recycled back from the medial to early-Golgi by the Vps74-COPI pathway, and any Ers1 that escapes recycling is readily targeted for degradation by the Dsc E3 ubiquitin ligases complex that maintains Golgi protein quality control. The Dsc complex surveils the Golgi, endosome, and vacuole, preventing accumulation of mislocalized proteins (Schmidt et al., 2019). This is reminiscent of sequential monitoring by protein quality control mechanisms in the endocytic pathway to clear mislocalized proteins and ensure proper organelle composition (Sardana and Emr, 2021; Sardana et al., 2019).

Maintenance of Ers1 at the early Golgi functionally contextualizes the observed suppression of the Golgi cargo recycling adaptor Erd1 (Hardwick and Pelham, 1990; Sardana et al., 2021). Loss of Ers1 alone does not cause Golgi cargo recycling defects. We speculate that suppression of the *erd* phenotype is due to modulation of the Golgi redox environment upon Ers1 overexpression. Alternatively, Ers1 may serve a redundant adaptor-like role for specific Golgi-resident proteins or facilitate the recruitment of cytosolic effectors to the Golgi. Indeed, a receptor-like role has been reported for the lysosomal cationic amino acid transporter and PQ-loop family protein PQLC2 (Talaia et al., 2021). Interaction between PQLC2 and WDR41 recruits the C9orf72–SMCR8–WDR41 complex to lysosomes, triggering downstream signaling. Interestingly, a recent study also proposed that cystinosin functions as a trafficking receptor for NHE3 (Khare et al., 2025). Additionally, other trafficking related defects such as the mis-localization of the chaperone-mediated autophagy receptor LAMP2A, the small GTPase Rab11, and Rab7 effector RILP have been proposed upon the loss of cystinosin (Zhang et al., 2017).

### A conserved role for Ers1 in redox homeostasis independent of cystine transport?

Both our data and prior studies demonstrate that Ers1 deletion does not result in cystine accumulation in yeast (Khare et al., 2025; Simpkins et al., 2016). On the other hand, loss of cystinosin function in mammalian cells is readily associated with intra-lysosomal cystine accumulation (Town et al., 1998). While lack of cystine accumulation could be due to redundant function by another transporter in yeast, our work provides a more plausible explanation for these observations. Intra-lysosomal cystine likely arises from proteolysis by lysosomal proteases. Cystinosin, which localizes to the lysosomal membrane, utilizes the proton gradient to transport cystine to the cytosol, where it is readily reduced and used for new protein synthesis. Unlike the lysosomes, protein hydrolysis or cystine generation is not expected to occur in the Golgi lumen. Given that Ers1 localizes to the early-Golgi at steady state, it is therefore unlikely to transport cystine across the Golgi membrane i*n vivo.* Not surprisingly, we did not observe cystine accumulation in *ers1*Δ mutant in total cells or in purified intact Golgi compartments. The absence of cystine-dependent transport by Ers1 in *Xenopus* oocytes, and function of putative cystine-binding Ers1 mutants in the *grx6Δ grx7Δ* temperature sensitivity assay, may also reflect a fundamental difference in solute preference between Ers1 and cystinosin. Future work is needed to shed light on whether Ers1 and cystinosin-LKG can also transport a solute that is structurally similar to cystine at the extra-lysosomal locations.

A role for Ers1 in redox homeostasis is supported by its genetic interaction with the ER and early Golgi-localized putative glutaredoxins Grx6 and Grx7. Glutaredoxins are thought to use glutathione (GSH) as the electron donor to reduce protein oxidation. Modulation of *grx6Δ grx7Δ* and *ero1-1* mutants in opposite directions by Ers1 suggests Ers1 activity contributes to a higher oxidized state in the ER/Golgi lumen. Interestingly, deletion of the C-terminal cytosolic tail disrupted Ers1’s ability to suppress *grx6Δ grx7Δ* mutant. Ers1 may transport an as-yet-unidentified solute across the Golgi membrane via residues distinct from those required for cystine transport, or it may perform a non-transport structural function by recruitment of an interactor in part via its cytosolic C-terminal tail.

Disruption of redox and glutathione homeostasis in cystinosis models has been reported extensively. Altered cytosolic cystine and cysteine levels likely drive this phenotype, particularly given that glutathione synthesis depends on cysteine availability. Our data further suggest that interactions with glutathione and redox homeostasis machinery warrant investigation in human cells. Notably, CTNS-LKG was able to complement the function of Ers1 in the *erd1*Δ and *grx6Δ grx7Δ* assays. CTNS and CTNS-LKG differ from each other in the length and composition of their cytosolic tails; CTNS-LKG lacks the C-terminal lysosomal targeting tyrosine motif, resulting in a substantial shift in cellular distribution, with reduced lysosomal and endosomal localization and increased cytoplasmic vesicles, Golgi, and plasma membrane localization. Although both isoforms can transport cystine, the ability of CTNS-LKG to complement Ers1 function raises the possibility that protein localization and interactions contribute to its function.

Our findings demonstrate that cystinosin’s role extends beyond lysosomal cystine transport and expand the conceptual framework for studying cystinosin and other PQ family proteins. There are several questions that will need to be addressed and will require future investigations. What solute, if any, does Ers1 transport across the Golgi membrane? Are CTNS and CTNS-LKG also capable of transporting solutes other than cystine? What roles does CTNS-LKG play in the extra-lysosomal locations? What is the role played by the C-terminal tail of Ers1? How does Ers1 crosstalk with Grx6 and Grx7? Importantly, is the role played by Ers1 in the early secretory pathway important for cystinosis pathology? If conserved in humans, these findings may open new therapeutic opportunities for cystinosis.

## Methods

### Fluorescence microscopy

Yeast cells were grown in synthetic dropout media to mid-log phase at 30° C and imaged at room temperature on glass coverslips or in glass-bottomed dishes. Images unless mentioned otherwise were captured using an ECLIPSE Ti2 inverted microscope, equipped with a 100x oil objective. Image acquisition and deconvolution were performed using the provided NIS Elements software. For CMAC staining of the vacuole, cells were treated with 100uM CMAC for 15min before imaging.

Images for Golgi puncta quantification were captured with a CSU-X spinning-disk confocal microscope system (Intelligent Imaging Innovations) using a DMI6000 microscope (Leica Microsystems) outfitted with a CSU-X1 spinning-disk confocal unit (Yokogawa Electric Corporation) with a QuantEM 512SC (Photometrics). The objective was a 100 × 1.46 NA Plan Apochromat oil immersion lens (Leica Microsystems). Quantification of Golgi puncta was implemented in Python. Briefly, a median filter was used to correct images for hot pixels, and a gaussian background subtraction was applied to remove uneven illumination. Puncta in the two fluorescent channels were segmented by thresholding using the Otsu method. Object based co-localization was used to compute the overlap ratio between the two channels for ∼250 puncta per dataset. Each puncta’s quantification data was plotted, with the median and confidence intervals annotated in violin plots.

For cellular localization of YFP tagged CTNS and Ers1 in Xenopus oocytes, the YFP chimeras was imaged on a confocal laser-scanning microscope (TCS SP5, Leica). A plasma membrane stain (CellMask™ Plasma Membrane Stains, Deep Red C10064, Thermo Fisher Scientific, USA) was used as a marker for PM co-localization. YFP fluorescence was excited with the 514 nm line (Argon laser gain: 40%) and the emission signal was collected between 524 to 566 nm (PMT gain: 115). The Deep Red stain was excited with the 594 nm line (Helium Neon laser gain: 35%) and the emission signal was collected between 614 and 708 nm (PMT gain: 26).

For immunofluorescence, CTNS-GFP and CTNS-LKG-GFP Knock-in HEK293 cells were grown on sterile glass coverslips in 6-well plates and fixed with 4% paraformaldehyde in PBS for 15 minutes at room temperature, permeabilized with 0.1% Triton X-100 in PBS for 5 minutes and blocked with 2% FBS in PBS for 1 hour at room temperature. Cells were incubated with primary antibodies (anti-GFP, anti-LAMP1, anti-GM130, anti-EEA1, and anti-EGFR) diluted (1:100) in blocking buffer for 3 hours at room temperature in a humid chamber. After washing, cells were incubated with species-specific Alexa Fluor–conjugated secondary antibodies (488 or 568) for 1 hour at room temperature in the dark. Coverslips were mounted with ProLong Gold Antifade Mountant with DAPI (Thermo Fisher Scientific) and imaged using a Nikon Ti2 confocal microscope. Images were processed using Nikon Elements and FIJI (ImageJ).

### Protein immunoprecipitation and biochemistry

Yeast lysates for immunoblotting were prepared from 2-5 OD600 equivalents of cells. Briefly, cells were harvested and incubated on ice for 1 h in 10 % trichloroacetic acid (TCA). After pelleting at 13G at 4C for 3min, cells were washed with cold acetone and lysed by beating with glass beads in 75 µl of 2 x urea buffer (50 mM Tris-HCl pH 7.5, 8 M urea, 2 % SDS, 1 mM ethylenediaminetetraacetic acid [EDTA]) for 5 min at room temperature, followed by incubation at 42 °C for 10 min. 75 µl of 2 x sample buffer (150 mM Tris HCl, pH 6.8, 8 M urea, 8 % SDS, 24 % glycerol, bromophenol blue, 100 mM dithiothreitol (DTT)) was added, and samples were vortexed for 5 min and again incubated at 42 °C for 10 min. After centrifugation, 0.2 OD equivalents of the cleared lysate were loaded on eight or 10% SDS-Polyacrylamide gels and transferred to nitrocellulose membranes (0.45 µm, GE healthcare) at 4 °C via wet transfer in transfer buffer (25 mM Tris, 192 mM Glycine, 10% v/v methanol, 0.006 % SDS) at 100 V for 90 min or semi-dry transfer in transfer buffer (300 mM Tris, 300 mM Glycine, 20% v/v ethanol, 0.1 % SDS) prior to immunoblotting.

The following antibodies and dilutions were used for western blotting: rabbit polyclonal anti-GFP (1:5000) (TP401; Torrey Pines Biolabs, Secaucus, NJ), mouse anti-alpha-Tubulin Hybridoma supernatant (1:3000) (clone 12G10, Developmental Studies Hybridoma Bank), rabbit anti-HA (1:3000) (GTX115044; Genetex), mouse monoclonal anti-FLAG (1:500) (012-22384; Wako), rabbit anti-Tul1, Dsc2, Dsc3, Dsc3, Ubx3 (1:2,000, Espenshade Lab), IRDye800 goat anti-rabbit (1:20,000) (926–32211; LI-COR Biosciences, Lincoln, NE) and IRdye680 goat anti-mouse (1:20,000) (926-32220; LI-COR Biosciences).

For immunoprecipitation of Ers1-GFP, cells were grown to OD600 0.8-1.0 in 250 ml of YPD and harvested on ice. Cells were washed and resuspended in Lysis Buffer (50 mM HEPES pH 7.5, 100mM NaCl, 1 mM EDTA pH 8.0, 10% glycerol, 0.005% NaN3, 1 mM PMSF, 1 X Nacalai Protease Inhibitor Tablet/50 ml). Cell extracts were prepared by glass bead beating for five cycles of 1 min vortexing with 2 min breaks on ice. Membranes were solubilized by nutating for 1h at 4 °C after addition of DDM to 1 % final concentration. Crude extracts were clarified and the lysates were precleared with 50 µl of sepharose beads and then incubated with 50 µl of GFP-nanobody resin at 4 °C for 4 hr. The resin was washed three times with Lysis buffer and then three times with HEPES-NP40 buffer (50mM HEPES pH7.5, 100 mM NaCl, 1mM EDTA, 0.2% NP40, 1mM PMSF, 1 X Nacalai Protease Inhibitor Tablet/50 ml). Bound proteins were eluted by addition of 100 µl elution buffer sample buffer (100 mM HEPES pH7.5, 1 % SDS), followed by incubation at 65 °C for 20 min.

### Proteomics sample preparation and analysis

For proteomics, the samples were reduced with 200 mM TCEP (Thermo, 77720) incubated at 55°C for 1 hour, then alkylated with 375 mM iodoacetamide incubated at room temperature for 30 minutes while protected from light. A precipitation solution (50% acetone, 49.9% ethanol, 0.1% acetic acid) was used to precipitate the proteins and remove SDS from the buffer. Protein pellets were resuspended in 100 mM TEAB (Thermo, 90114), then digested with 250 ng of Trypsin Gold (Promega, V5280) while nutating overnight at 37°C. The samples were acidified with 10% FA to quench the digestion, then desalted with homemade C18 solid-phase extraction columns. Briefly, C18 sorbent was extracted from a Sep-Pak 200 mg C18 cartridge (Waters, WAT054945) and distributed among empty micro-spin columns (Thermo, 89879). The spin columns containing the C18 sorbent were washed with 80% acetonitrile in 0.1% acetic acid and equilibrated with 0.1% TFA. The peptide samples were added to the spin column, washed with 0.1% acetic acid, then eluted with 80% acetonitrile in 0.1% acetic acid and dried in a Speed-Vac. The dried peptide samples were dissolved in 0.1% TFA and analyzed with a timsTOF HT mass spectrometer (Bruker) coupled with a NanoElute2 reverse phase liquid chromatography system (Bruker). A two-column separation method was employed for the LC including a 5 mm PepMap Neo Trap Cartridge (Thermo, 174500) and a homemade 10 cm by 100 µm analytical column packed with 1.9 µm C18 beads (Dr. Maisch, r119.aq.0001). A 25 minute elution gradient for peptide separation consisted of solvent A (0.1% formic acid in water) and steadily increasing concentration of solvent B (0.1% formic acid in acetonitrile) from 2% to 35% B. Mass spectrometric data was obtained in data-independent acquisition mode with parallel accumulation-serial fragmentation (diaPASEF), using 12 cycles with estimated cycle time of 1.7 seconds. Thirty-four DIA windows of 25 Da width and 2 mobility windows each were acquired with a ramp time of 75 ms and covered a mass range of 350 to 1250 Da and a mobility range of 0.64 1/K0 [Vs/cm-2] to 1.37 1/K0 [Vs/cm-2]. Collision energy was programmed to increase as a function of ion mobility, from 20 eV at 0.6 1/K0 [Vs/cm-2] to 59 eV at 1.6 1/K0 [Vs/cm-2]. Prior to running the samples, the elution voltage was calibrated for 1/K0 ratios using three ions (m/z 622, 922, 1222) from the ESI-L Tuning Mix (Agilent, G1969-85000) with the timsControl software (Bruker).

The data was analyzed using DIA-NN (v1.8) (Demichev et al., 2019). The diaPASEF runs were searched against an in silico spectral library that was generated using the same runs and a yeast proteome FASTA digest. Search parameters included the following: protease was set to ‘Trypsin/P’ with 3 missed cleavages allowed, 2 variable modifications maximum, cysteine carbamidomethylation and oxidized methionine. Peptide length range was set from 7-30 residues with precursor charge range 1-4 in an m/z range of 300-1800 and a fragment ion m/z range from 100-1800. Precursors were filtered for 1% FDR with MBR, double-pass neural network classifier, high accuracy LC quantification, and retention time-dependent cross-run normalization.

Downstream differential expression analysis was performed on the protein-level quantitative DIA-NN outputs using a customized linear modeling framework called MAGMA based on the Limma (Ritchie et al., 2015) and MSstatsTMT (Huang et al., 2020) R packages, consistent with approaches previously described for differential analysis of mass-spectrometry proteomics data. Briefly, protein abundance differences between experimental conditions were quantified by estimating fold changes and associated statistical significance using moderated linear models. The core analysis framework and statistical modeling strategy have been previously described in published work (Zhou et al., 2022), with internal extensions applied here to support DIA-based quantification.

For DIA-specific handling of missing values, imputation was applied under a missing-not-at-random assumption. Specifically, for proteins missing across all replicates of a given condition, values were imputed using the minimum observed intensity for that protein within the corresponding condition replicate. This approach assumes left-censoring due to limits of detection and leverages the increased quantitative consistency of DIA data, where systematic absence across replicates is interpreted as biologically meaningful signal loss rather than stochastic dropout. Proteins were considered statistically significant if they exhibited fold change greater than 2 and a Benjamini–Hochberg–adjusted false discovery rate (FDR) < 0.05.

### Golgi immunoisolation and protease protection assay

Yeast cells were grown to 0.8-1.0 OD in 50ml YPD and 40OD was collected. Cells were spheroplasted with zymolyase, then resuspended in lysis buffer (50mM HEPES/KOH pH7.4, 50mM potassium acetate, 0.2 M sorbitol, 2 mM EDTA, 20mM NaF, 20mM NaN3, 1mM PMSF, 1 X Roche cOmplete Protease Inhibitor Tablet/50 ml) and lysed with 10 strokes using a Dounce homogenizer. The lysate was collected after clarification at 1G 2min and incubated with 40ul of GFP-Nb magnetic beads (12.5% slurry) for 15min at 4C with rotating. The beads were washed three times, then snap-frozen in liquid nitrogen and stored at -80C until metabolomics analysis. For the protease protection assay, samples were treated with 0.1ug/ul Proteinase K (NEB 8107G) with or without 1% Triton-X 100 for 30 minutes on ice. Samples were then precipitated with 10% TCA, washed with cold acetone, resuspended in 2 x sample buffer (125 mM Tris HCl, pH 7.5, 5mM EDTA, 4 % SDS, 5 % glycerol, bromophenol blue, 100mM DTT) and boiled at 65C for 20min before resolving on a 10% SDS-PAGE gel and subjected to immunoblotting.

### Measurement of ER redox

Yeast strains transformed with a plasmid expressing ER-targeted roGFPiE were grown in low fluorescence SD-ura (without folic acid and riboflavin), harvested by centrifugation, washed once with water and then resuspended in 100 mM pH5.5 MES buffer. 4 OD of cells in 200 µl MES buffer were added to each well of a black 96-well flat bottom plate (Greiner) and measured with a Tecan plate reader. Excitation/emission wavelengths were 395/505 nm and 465/505 nm with a bandwidth of 10 nm.

### Yeast metabolite extraction and analysis

The extraction protocol was adapted and modified from a published work (Thirumalaikumar et al., 2021). Proteins and metabolites were extracted using a methyl tert-butyl ether/methanol/water solvent system, which separates molecules into pellets (proteins), organics (lipids), and an aqueous phase (primary and secondary metabolites). Equal volumes of the polar and lipid fractions were dried in a centrifugal evaporator and stored at −80 °C until further processing. The dried aqueous phase was measured using ultraperformance liquid chromatography coupled to an Exactive mass spectrometer (Thermo Fisher Scientific, Waltham, MA, USA) in positive and negative ionization modes, as described a published work (Thirumalaikumar et al., 2021). A 2 μL sample (the dried aqueous fraction was resuspended in 200 μL of UHPLC-grade water) was loaded per injection. The LC/MS data were processed using Expressionist Refiner MS 11.0 (Genedata AG, Basel, Switzerland). Settings were as follows: chromatogram alignment (RT search interval, 0.5 min), peak detection (summation window, 0.09 min; minimum peak size, 0.03 min; gap/peak ratio, 50%; smoothing window, 5 points; center computation by intensity-weighted method with threshold at 70%; and boundary determination using inflection points), isotope clustering (RT tolerance at 0.015 min, m/z tolerance 5 p.p.m., and allowed charges 1–5), filtering for a single peak not assigned to an isotope cluster, adduct detection, and cluster grouping (RT tolerance 0.05 min, m/z tolerance 5 p.p.m., and maximum intensity of side adduct 100 000%). All metabolite clusters were matched to in-house libraries of reference compounds, allowing a 10-p.p.m. mass and dynamic retention time deviation (maximum 0.1 min). Raw metabolites’ intensities were normalized to the median of chromatogram intensity in order to correct for variation in sample extraction and measurement. Prior statistical analysis data were subjected to log2 transformation. P-value was calculated using unpaired Student’s t-test with two-tailed distribution; n = 4 of biological replicates.

### Generation of Knock-In CTNS and CTNS-LKG-GFP cell lines

cDNAs encoding CTNS-GFP and CTNS-LKG-GFP were integrated at the endogenous CTNS locus in HEK293T cells using **C**RISPR-mediated, **H**omology-independent, **P**CR-product **I**ntegration (CHoP-In) approach for genome editing published previously (Manna et al., 2019). Cells were cultured in Dulbecco’s Modified Eagle Medium containing 4.5 mg/ml D-glucose and 2 mM L-glutamine supplemented with 10% fetal bovine serum at 37°C in a humidified incubator with 5% CO₂. Oligonucleotides encoding the sgRNA targeting exon 3 right after the start codon of CTNS were cloned in the BbsI sites of pX330-Cas9 (Addgene #42230). Primers for generating CHoP-In integration donors were designed to amplify the integration fragment containing a left homology arm followed by cDNA for CTNS or CTNS-LKG, C-terminal moxGFP, P2A site, puromycin resistance gene and a right homology arm. Cells were transfected using Avalanche Omni transfection reagent according to manufacturers’ instructions. Successfully edited cells were selected 48-72 hours post-transfection with 0.3 µg/ml puromycin, enriched by flow cytometry sorting, and subsequently cultured to generate monoclonal cell lines.

### Two-Electrode Voltage-Clamp electrophysiology in *Xenopus* oocytes

The coding sequence of CTNS- ΔGYDQL, CTNS-LKG, and Ers1 were amplified and cloned into Xenopus oocyte expression vectors (with and without C-terminus YFP) by the advanced uracil excision-based cloning technique described previously (Li et al., 2020). After DNA linearization with NheI, cRNA was synthesized using the New England Biolabs T7 in vitro transcription kit following the manufacturer’s guidelines.

For protein expression in oocytes, stage V and VI oocytes were harvested from Xenopus laevis and maintained in ND96 solution (96 mM NaCl, 2 mM KCl, 1.8 mM CaCl2, 1 mM MgCl2, 2.5 mM Na Pyruvate, 5 mM Hepes – pH 7.5, 50 mg mL− 1 gentamycin and 0.4 g L− 1 BSA). Oocytes were defolliculated by collagenase treatment in ND96 without CaCl2, gentamycin and BSA and maintained overnight at 18 °C in complete ND96 solution as described initially. All animal procedures, including husbandry, oocyte harvesting and post-treatment, were performed in accordance with Cornell University IACUC Protocol number 2017–0139. Oocytes were microinjected with 50 nl of cRNA (500 ng/μl) using an oil-driven injection system (Nanoject II Auto-Nanoliter Injector, Drummond Scientific Company, US) and then incubated for 3d at 18 °C in complete ND96 solution. Whole-cell currents were measured using conventional Two-Electrode Voltage-Clamp technique (Axoclamp 900A amplifier and Digidata 1550B-PClamp 1 data acquisition system, Molecular Devices) while under continuous perfusion of bath solutions containing 96 mM NaCl, 1 mM KCl, 2 mM MgCl2, 1.8 mM CaCl2, buffered with 10 mM MES at pH 5.5 or pH 6.5, with the osmolarity adjusted to 220 mOsmol/kg with sorbitol), and with or without the addition of 1 mM cystine, cysteine, GSH, or GSSG.

## Acknowledgements

We thank Drs. Scott Emr, Chris Fromme, Carolyn Sevier, and Martin Graef for feedback on the manuscript. We thank Dr. Teresa Dunn for FLAG-Orm2 plasmid, and Dr. Peter Espenshade for sera against yeast Dsc complex components (Tul1, Dsc2, Dsc3, Ubx3). We thank Dr. Carolyn Sevier for *ero1-1* yeast strain and the ER-roGFP-iE plasmid. We thank Dr. Scott Emr for the Tul1 overexpression plasmid. We thank Dr. Feng Zhang for the pX330-U6-Chimeric_BB-CBh-hSpCas9 plasmid (Addgene# 42230), and Dr. Laurence Pelletier for the qTAG-C-moxGFP-Puro plasmid (Addgene# 207721). This work was supported by start-up funds from Cornell University College of Veterinary Medicine to RS, Cornell University CVM graduate student fellowship to JZ, and National Institute of General Medical Sciences of the National Institutes of Health award numbers 5R35GM150863 to RS, and R35GM153298 to AS.

---

**Figure S1.**
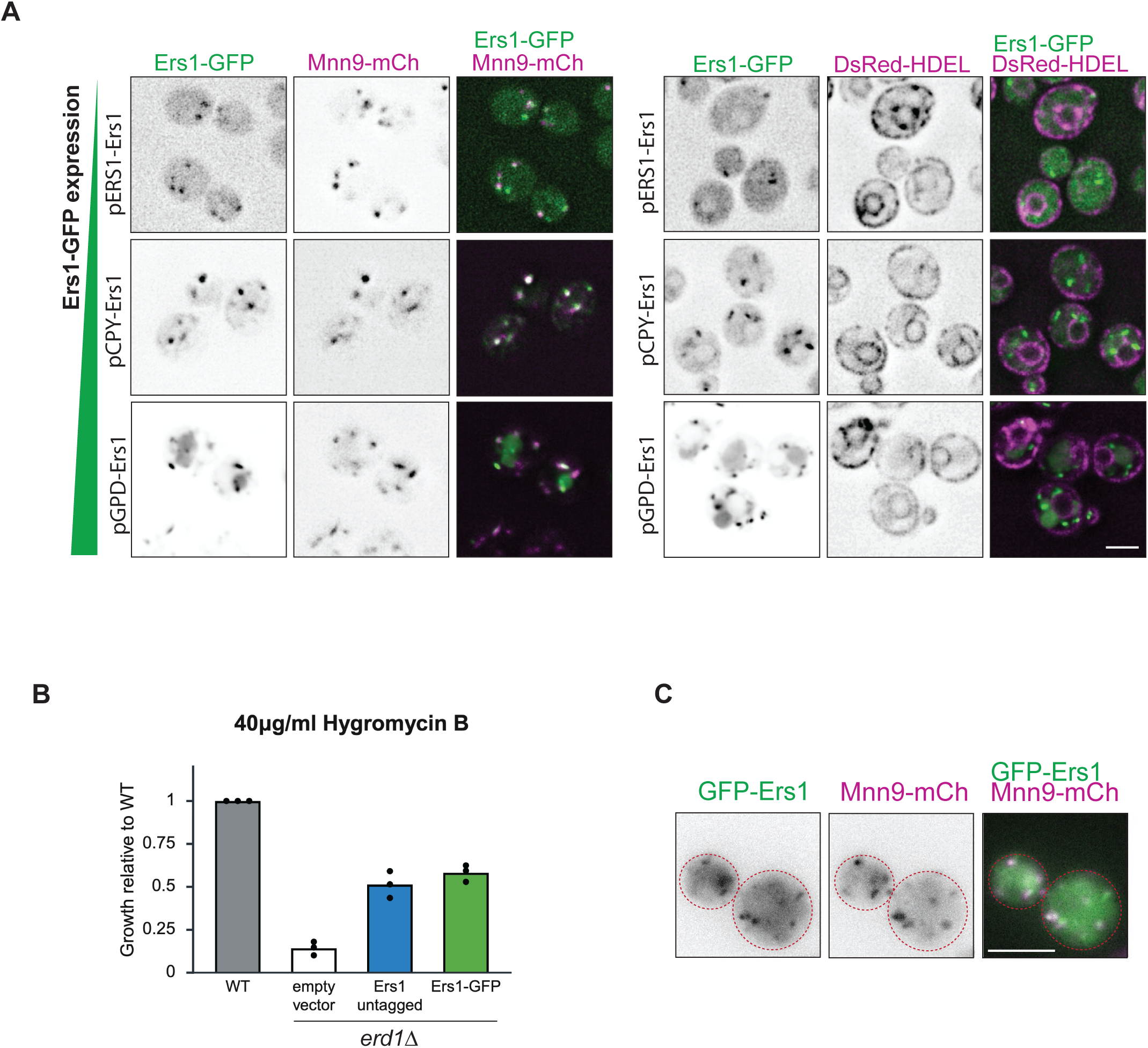
(A) Colocalization of C terminally tagged Ers1-GFP expressed under the control of the ERS1 promoter, CPY promoter, or GPD promoter with either an early Golgi marker (Mnn9-mCherry) or ER marker (DsRed-HDEL). (B) Growth of WT and *erd1*Δ cells in liquid culture with 40ug/ml Hygromycin B. A plasmid encoding for untagged or C-terminally GFP-tagged Ers1 under the control of the GPD promoter was transformed into *erd1*Δ cells. Bar graphs represent averages from three independent experiments, shown as individual dots. (C) Colocalization of N terminally tagged GFP-Ers1 expressed under the control of the CPY promoter with an early Golgi marker (Mnn9-mCherry). Scale bar 5 µm.

**Figure S2.**
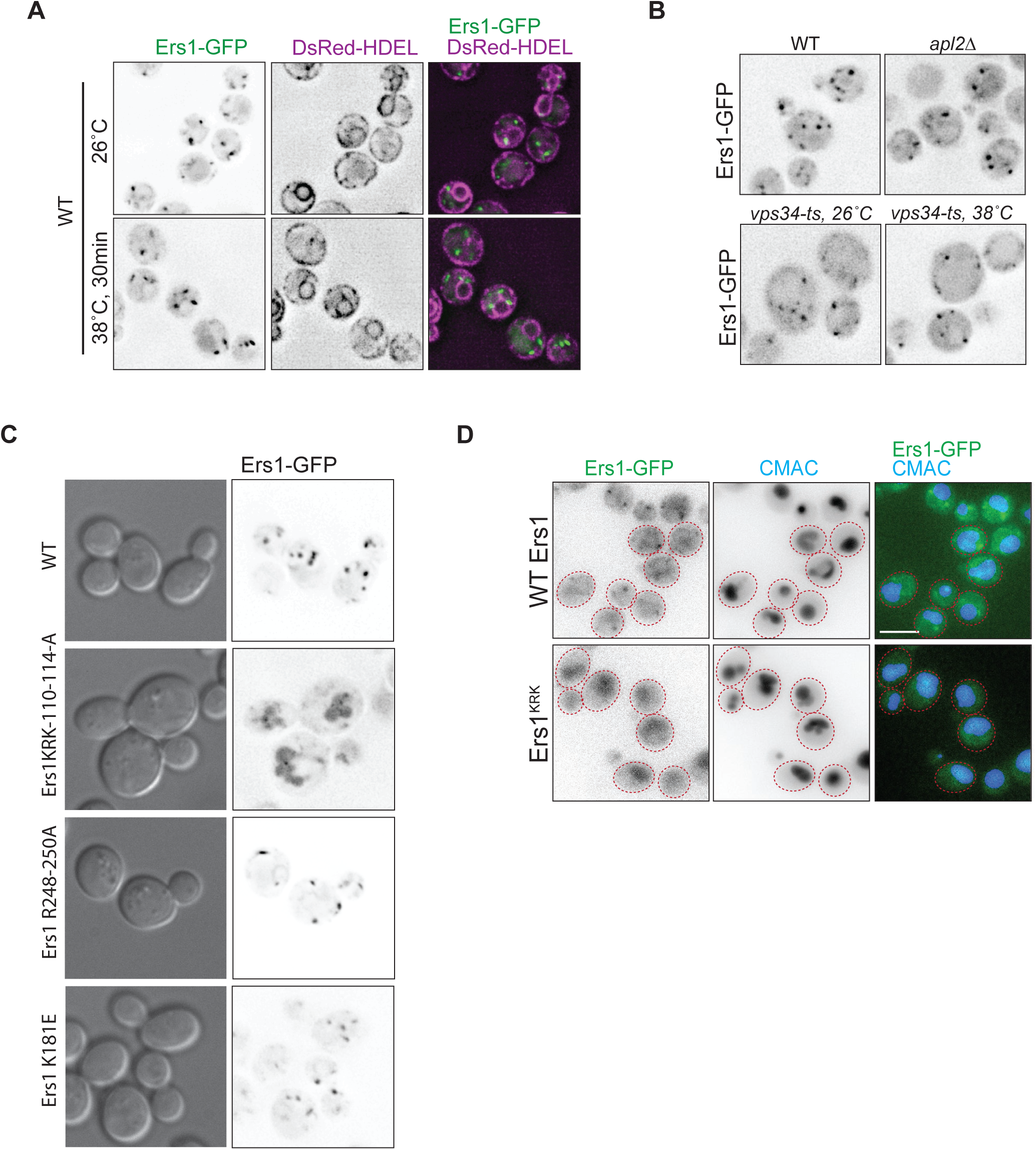
(A) Colocalization of Ers1-GFP in wild-type cells with an ER marker (DsRed-HDEL). Cells were grown to mid-log phase at 26°C and then incubated at 38°C for 30 min before imaging. (B) Localization of Ers1-GFP in the AP-1 mutant *apl2*Δ and PI3K mutant *vps34-ts*. The *vps34-ts* mutant was grown to mid-log phase at 26°C and then incubated at 38°C for 30 min before imaging. (C) Localization of GFP-tagged wild-type and basic patch mutants of Ers1- (K110A, R113A, K114A) (R248A R250A) (K181E) in wild-type cells. (D) Localization of wild-type and mutant (K110A, R113A, K114A) Ers1-GFP in wild-type cells. Cells were grown in a synthetic medium and vacuoles were stained with CMAC.

**Figure S4.**
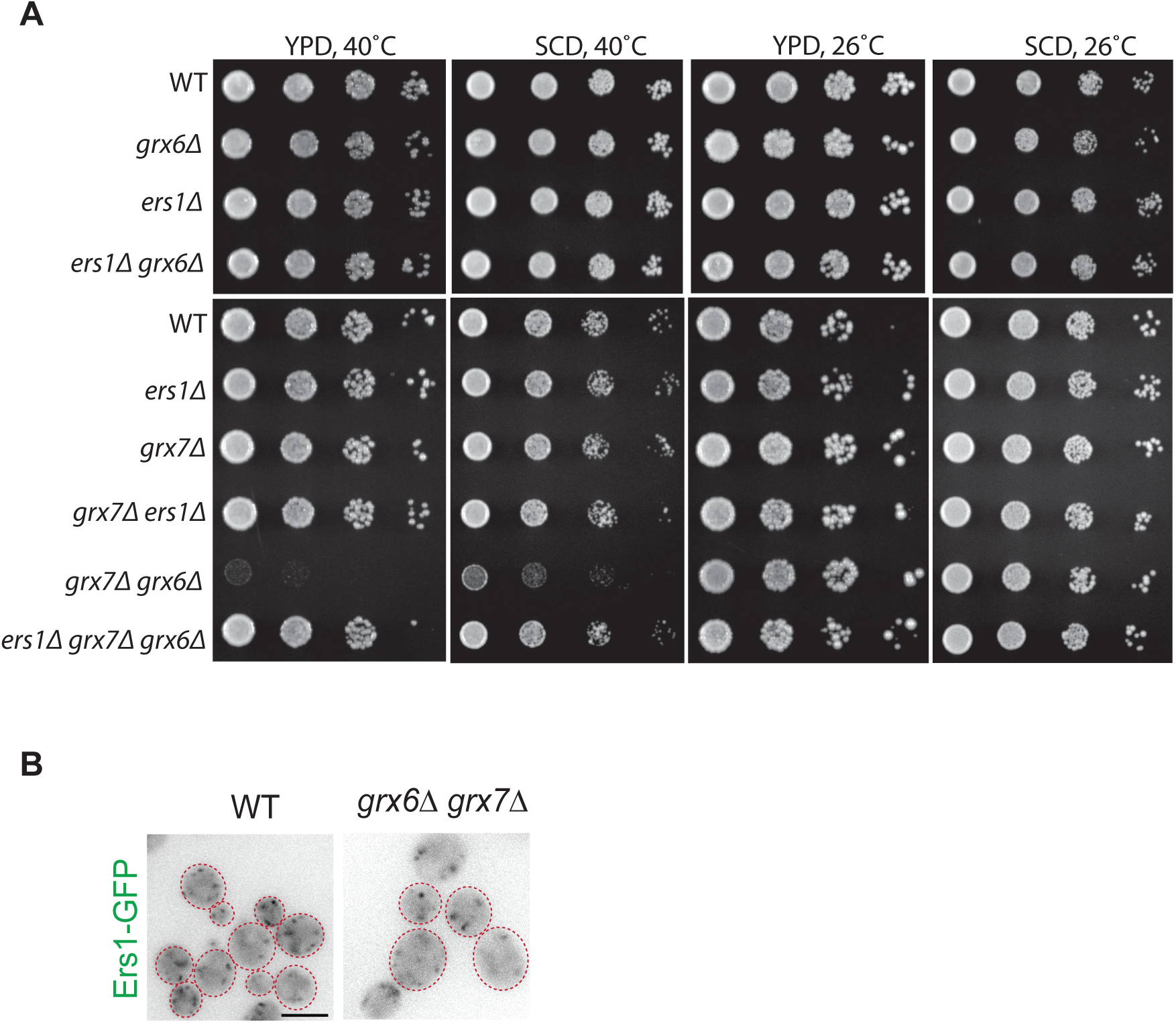
(A) Growth under temperature stress of wild-type, *ers1Δ, grx6Δ, ers1Δgrx6Δ, grx7Δ, grx7Δ ers1Δ, grx6Δgrx7Δ*, and *grx6Δgrx7Δers1Δ* cells. Cells were spotted as seriated dilutions on YPD or SCD plates and incubated at 26°C or 40°C for 3 days. (B) Localization of Ers1-GFP in wild-type and *grx6Δgrx7Δ* cells. Cells were grown in a synthetic medium at 30°C to mid-log phase.

**Figure S5.**
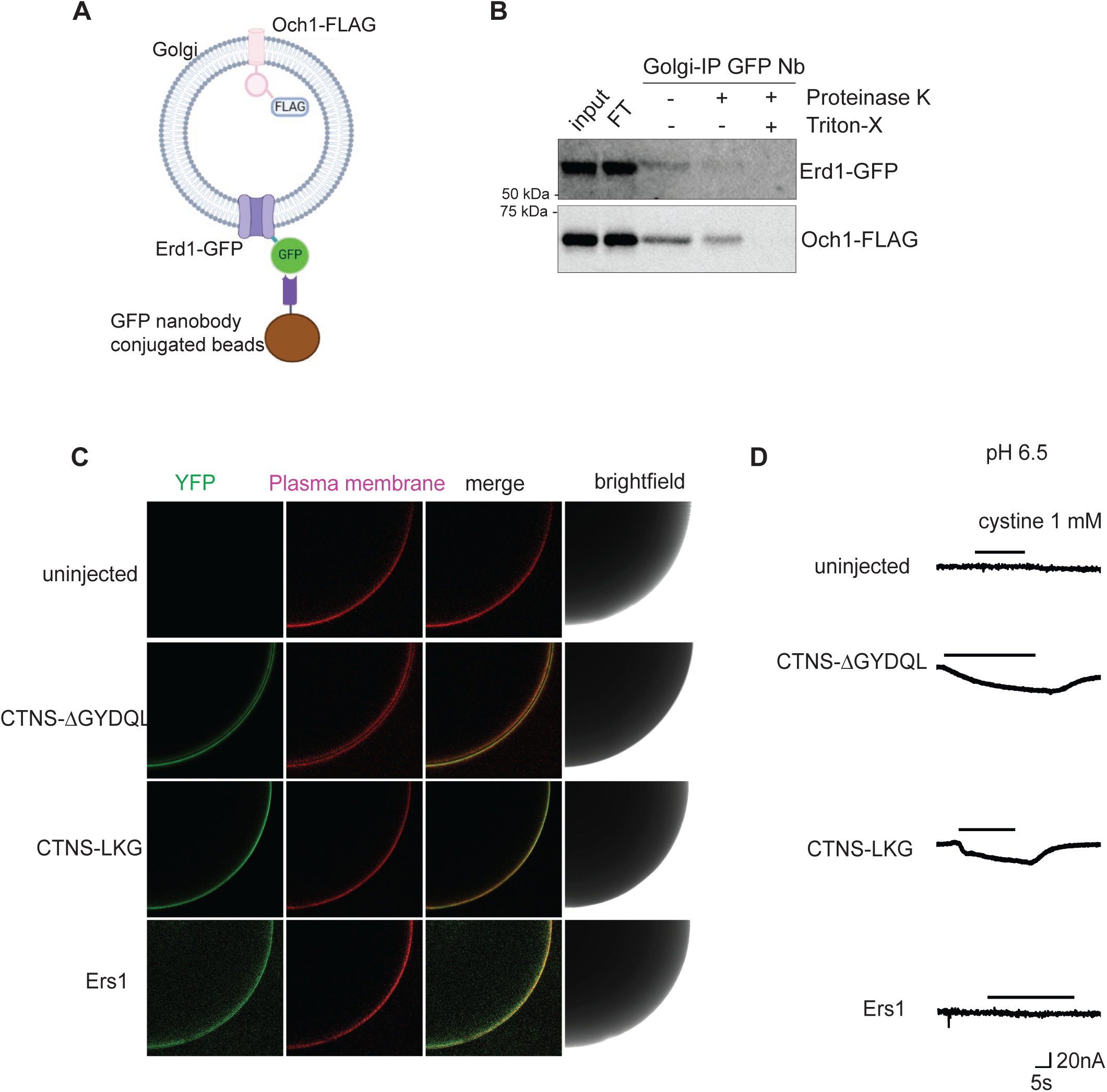
(A) Schematic for the strategy for Golgi IP. GFP nanobody conjugated magnetic beads are used for immunoisolation of early-Golgi localized Erd1-GFP. Lumen facing FLAG tag on early-Golgi localized Och1 is used for confirmation of Golgi intactness by proteinase protection assay. (B) Western blot for Erd1-GFP and Och1-FLAG in immunoisolated Golgi vesicles treated with proteinase K and Triton-X. (C) Localization of YFP-tagged CTNS-ΔGYDQL, CTNS-LKG, and Ers1 in oocytes and colocalization with the plasma membrane marker Deep Red. (D) Representative currents induced by addition of 1mM cystine at pH 6.5 in uninjected oocytes and oocytes expressing CTNS-ΔGYDQL, CTNS-LKG, and Ers1.

**Figure S6.**
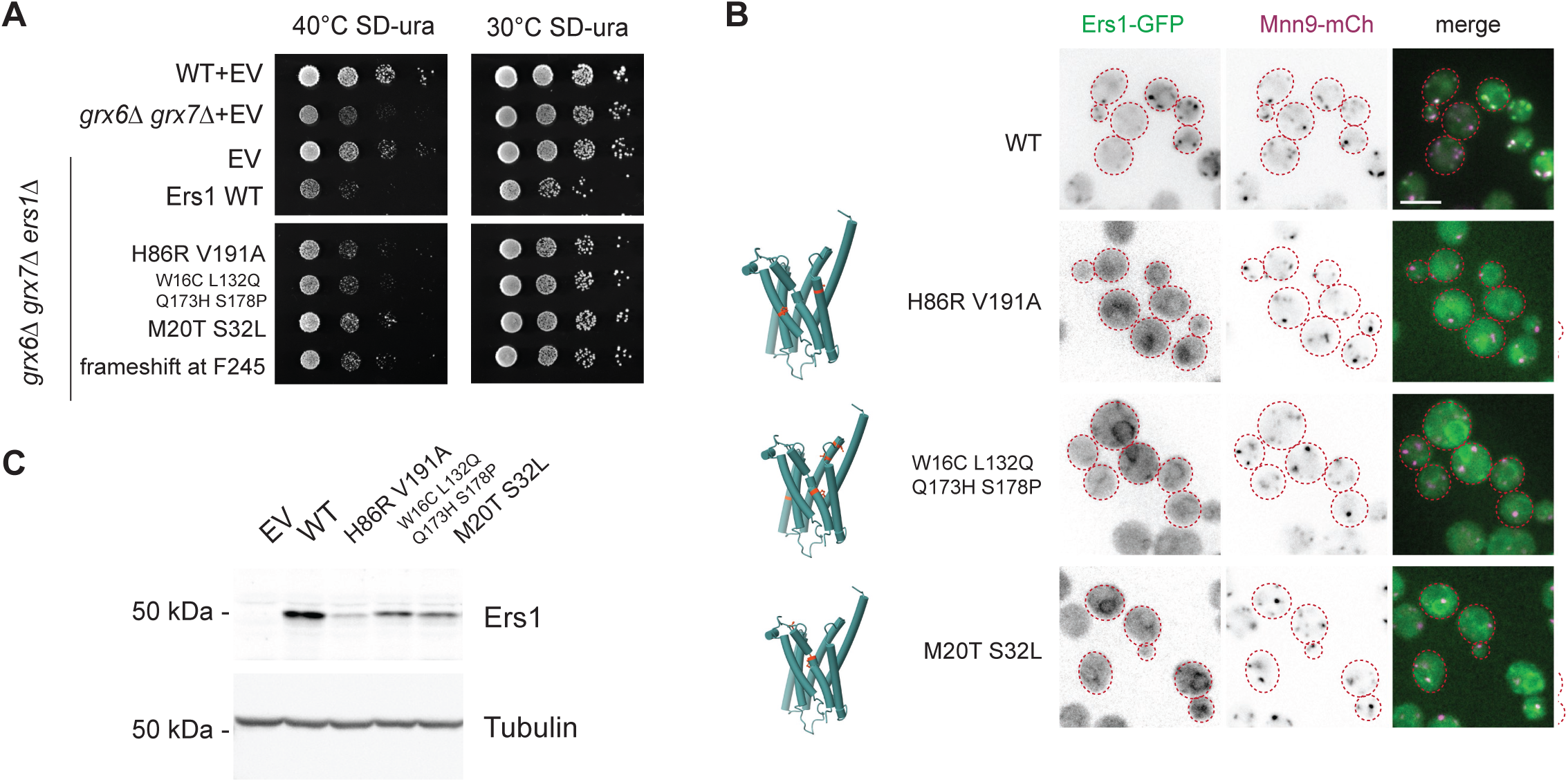
(A) Growth under temperature stress of wild-type, *grx6Δgrx7Δ*, and *grx6Δgrx7Δers1Δ* cells expressing wild-type Ers1 or Ers1 missense mutants identified in the random mutagenesis screen. Cells were spotted as seriated dilutions on SD-ura plates and incubated at 30°C or 40°C for 3 days. (B) Fluorescence microscopy analysis of GFP-tagged Ers1 missense mutants identified in the random mutagenesis screen with an early Golgi marker (Mnn9-mCherry). (C) Western blot analysis of whole cell lysates from cells expressing the GFP-tagged Ers1 missense mutants identified in the random mutagenesis screen.

## References

Anand, M., Rush, J.S., Ray, S., Doucey, M.A., Weik, J., Ware, F.E., Hofsteenge, J., Waechter, C.J., Lehrman, M.A., 2001. Requirement of the Lec35 gene for all known classes of monosaccharide-P-dolichol-dependent glycosyltransferase reactions in mammals. Mol. Biol. Cell 12, 487–501. 10.1091/mbc.12.2.487

Andrzejewska, Z., Nevo, N., Thomas, L., Chhuon, C., Bailleux, A., Chauvet, V., Courtoy, P.J., Chol, M., Chiara Guerrera, I., Antignac, C., 2016. Cystinosin is a Component of the Vacuolar H+-ATPase-Ragulator-Rag Complex Controlling Mammalian Target of Rapamycin Complex 1 Signaling. J. Am. Soc. Nephrol. 27, 1678–1688. 10.1681/ASN.2014090937

Bellomo, F., Taranta, A., Petrini, S., Venditti, R., Rocchetti, M.T., Rega, L.R., Corallini, S., Gesualdo, L., de Matteis, M.A., Emma, F., 2016. Carboxyl-Terminal SSLKG Motif of the Human Cystinosin-LKG Plays an Important Role in Plasma Membrane Sorting. PLoS One 11. 10.1371/journal.pone.0154805

Berquez, M., Chen, Z., Festa, B.P., Krohn, P., Keller, S.A., Parolo, S., Korzinkin, M., Gaponova, A., Laczko, E., Domenici, E., Devuyst, O., Luciani, A., 2023. Lysosomal cystine export regulates mTORC1 signaling to guide kidney epithelial cell fate specialization. Nat. Commun. 14. 10.1038/s41467-023-39261-3

Besouw, M., Masereeuw, R., Van Den Heuvel, L., Levtchenko, E., 2013. Cysteamine: An old drug with new potential. Drug Discov. Today 18, 785–792. 10.1016/j.drudis.2013.02.003

Birk, J., Meyer, M., Aller, I., Hansen, H.G., Odermatt, A., Dick, T.P., Meyer, A.J., Appenzeller-Herzog, C., 2013. Endoplasmic reticulum: reduced and oxidized glutathione revisited. J. Cell Sci. 126, 1604–1617. 10.1242/jcs.117218

Bräuer, P., Parker, J.L., Gerondopoulos, A., Zimmermann, I., Seeger, M.A., Barr, F.A., Newstead, S., 2019. Structural basis for pH-dependent retrieval of ER proteins from the Golgi by the KDEL receptor. Science 363, 1103–1107. 10.1126/science.aaw2859

Capitani, M., Sallese, M., 2009. The KDEL receptor: new functions for an old protein. FEBS Lett. 583, 3863–3871. 10.1016/j.febslet.2009.10.053

Carneiro, A., Jones, D.H., 2026. Advances in Pharmacological Treatments for Cystinosis: Cysteamine and Its Alternatives. ACS Pharmacol. Transl. Sci. 10.1021/acsptsci.5c00633

Cela, I., Dufrusine, B., Rossi, C., Luini, A., De Laurenzi, V., Federici, L., Sallese, M., 2022. KDEL Receptors: Pathophysiological Functions, Therapeutic Options, and Biotechnological Opportunities. Biomedicines 10. 10.3390/biomedicines10061234

Cherqui, S., 2012. Cysteamine therapy: A treatment for cystinosis, not a cure. Kidney Int. 81, 127–129. 10.1038/ki.2011.301

Cherqui, S., Courtoy, P.J., 2017. The renal Fanconi syndrome in cystinosis: pathogenic insights and therapeutic perspectives. Nat. Rev. Nephrol. 13, 115. 10.1038/NRNEPH.2016.182

Chol, M., Nevo, N., Cherqui, S., Antignac, C., Rustin, P., 2004. Glutathione precursors replenish decreased glutathione pool in cystinotic cell lines. Biochem. Biophys. Res. Commun. 324, 231–235. 10.1016/J.BBRC.2004.09.033

David, D., Princiero Berlingerio, S., Elmonem, M.A., Oliveira Arcolino, F., Soliman, N., Van Den Heuvel, B., Gijsbers, R., Levtchenko, E., 2019. Molecular Basis of Cystinosis: Geographic Distribution, Functional Consequences of Mutations in the CTNS Gene, and Potential for Repair. Nephron 141, 133–146. 10.1159/000495270

Demichev, V., Messner, C.B., Vernardis, S.I., Lilley, K.S., Ralser, M., 2019. DIA-NN: neural networks and interference correction enable deep proteome coverage in high throughput. Nature Methods 2019 17:1 17, 41–44. 10.1038/s41592-019-0638-x

Downes, K.W., Zanetti, G., 2025. Mechanisms of COPII coat assembly and cargo recognition in the secretory pathway. Nat. Rev. Mol. Cell Biol. 26, 910–925. 10.1038/s41580-025-00839-y

Ellgaard, L., Sevier, C.S., Bulleid, N.J., 2018. How Are Proteins Reduced in the Endoplasmic Reticulum? Trends Biochem. Sci. 43, 32–43. 10.1016/j.tibs.2017.10.006

Elmonem, M.A., Veys, K.R., Soliman, N.A., Van Dyck, M., Van Den Heuvel, L.P., Levtchenko, E., 2016. Cystinosis: a review. Orphanet J. Rare Dis. 11. 10.1186/S13023-016-0426-Y

Festa, B.P., Chen, Z., Berquez, M., Debaix, H., Tokonami, N., Prange, J.A., Hoek, G. Van De, Alessio, C., Raimondi, A., Nevo, N., Giles, R.H., Devuyst, O., Luciani, A., 2018. Impaired autophagy bridges lysosomal storage disease and epithelial dysfunction in the kidney. Nat. Commun. 9. 10.1038/s41467-017-02536-7

Frand, A.R., Kaiser, C.A., 1998. The ERO1 gene of yeast is required for oxidation of protein dithiols in the endoplasmic reticulum. Mol. Cell 1, 161–170. 10.1016/S1097-2765(00)80017-9

Gahl, W.A., Bashan, N., Tietze, F., Bernardini, I., Schulman, J.D., 1982. Cystine Transport Is Defective in Isolated Leukocyte Lysosomes from Patients with Cystinosis. Science (1979). 217, 1263–1265. 10.1126/SCIENCE.7112129

Gaide Chevronnay, H.P., Janssens, V., Van Der Smissen, P., Liao, X.H., Abid, Y., Nevo, N., Antignac, C., Refetoff, S., Cherqui, S., Pierreux, C.E., Courtoy, P.J., 2015. A mouse model suggests two mechanisms for thyroid alterations in infantile cystinosis: decreased thyroglobulin synthesis due to endoplasmic reticulum stress/unfolded protein response and impaired lysosomal processing. Endocrinology 156, 2349–2364. 10.1210/EN.2014-1672

Gao, X.D., Wang, J., Keppler-Ross, S., Dean, N., 2005. ERS1 encodes a functional homologue of the human lysosomal cystine transporter. FEBS J. 272, 2497–2511. 10.1111/J.1742-4658.2005.04670.X

Guo, X., Schmiege, P., Assafa, T.E., Wang, R., Xu, Y., Donnelly, L., Fine, M., Ni, X., Jiang, J., Millhauser, G., Feng, L., Li, X., 2022. Structure and mechanism of human cystine exporter cystinosin. Cell 185, 3739–3752.e18. 10.1016/J.CELL.2022.08.020

Hardwick, K.G., Pelham, H.R.B., 1990. ERS1 a seven transmembrane domain protein from Saccharomyces cerevisiae. Nucleic Acids Res. 18, 2177–2177. 10.1093/NAR/18.8.2177

Huang, T., Choi, M., Tzouros, M., Golling, S., Pandya, N.J., Banfai, B., Dunkley, T., Vitek, O., 2020. MSstatsTMT: Statistical Detection of Differentially Abundant Proteins in Experiments with Isobaric Labeling and Multiple Mixtures. Molecular and Cellular Proteomics 19, 1706–1723. 10.1074/mcp.RA120.002105

Hwang, C., Sinskey, A.J., Lodish, H.F., 1992. Oxidized redox state of glutathione in the endoplasmic reticulum. Science 257, 1496–1502. 10.1126/science.1523409

Izquierdo, A., Casas, C., Mühlenhoff, U., Lillig, C.H., Herrero, E., 2008. Saccharomyces cerevisiae Grx6 and Grx7 Are Monothiol Glutaredoxins Associated with the Early Secretory Pathway. Eukaryot. Cell 7, 1415–1426. 10.1128/EC.00133-08

Jain, A., Zoncu, R., 2021. Picking the arginine lock on PQLC2 cycling. Proc. Natl. Acad. Sci. U. S. A. 118. 10.1073/pnas.2112682118

Jamalpoor, A., Othman, A., Levtchenko, E.N., Masereeuw, R., Janssen, M.J., 2021. Molecular Mechanisms and Treatment Options of Nephropathic Cystinosis. Trends Mol. Med. 27, 673–686. 10.1016/J.MOLMED.2021.04.004

Jeźégou, A., Llinares, E., Anne, C., Kieffer-Jaquinod, S., O’Regan, S., Aupetit, J., Chabli, A., Sagné, C., Debacker, C., Chadefaux-Vekemans, B., Journet, A., Andreć, B., Gasnier, B., 2012. Heptahelical protein PQLC2 is a lysosomal cationic amino acid exporter underlying the action of cysteamine in cystinosis therapy. Proc. Natl. Acad. Sci. U. S. A. 109. 10.1073/pnas.1211198109

Johnson, J.L., Napolitano, G., Monfregola, J., Rocca, C.J., Cherqui, S., Catz, S.D., 2013. Upregulation of the Rab27a-dependent trafficking and secretory mechanisms improves lysosomal transport, alleviates endoplasmic reticulum stress, and reduces lysosome overload in cystinosis. Mol. Cell. Biol. 33, 2950–2962. 10.1128/MCB.00417-13

Kalatzis, V., Cherqui, S., Antignac, C., Gasnier, B., 2001a. Cystinosin, the protein defective in cystinosis, is a H(+)-driven lysosomal cystine transporter. EMBO J. 20, 5940–5949. 10.1093/EMBOJ/20.21.5940

Kalatzis, V., Cherqui, S., Antignac, C., Gasnier, B., 2001b. Cystinosin, the protein defective in cystinosis, is a H(+)-driven lysosomal cystine transporter. EMBO J. 20, 5940–5949. 10.1093/emboj/20.21.5940

Kellokumpu, S., 2019. Golgi pH, Ion and Redox Homeostasis: How Much Do They Really Matter? Front. Cell Dev. Biol. 7. 10.3389/fcell.2019.00093

Khare, V., Farre, J.C., Rocca, C., Kbaich, M.A., Tang, C., Ma, X., Biederman, K., Mathur, I., Grau, R.A.B.-, Sivakumar, A., Chen, R., Catz, S.D., Cherqui, S., 2025. Cystinosin is involved in Na+/H+ Exchanger 3 trafficking in the proximal tubular cells: new insights in the renal Fanconi syndrome in cystinosis. bioRxiv 2025.02.12.637793. 10.1101/2025.02.12.637793

Kriel, J., Haesendonckx, S., Rubio-Texeira, M., Van Zeebroeck, G., Thevelein, J.M., 2011. From transporter to transceptor: signaling from transporters provokes re-evaluation of complex trafficking and regulatory controls: endocytic internalization and intracellular trafficking of nutrient transceptors may, at least in part, be governed by the…. Bioessays 33, 870–879. 10.1002/bies.201100100

Kumar, C., Igbaria, A., D’Autreaux, B., Planson, A.G., Junot, C., Godat, E., Bachhawat, A.K., Delaunay-Moisan, A., Toledano, M.B., 2011. Glutathione revisited: a vital function in iron metabolism and ancillary role in thiol-redox control. EMBO J. 30, 2044–2056. 10.1038/emboj.2011.105

Letourneur, F., Gaynor, E.C., Hennecke, S., Démollière, C., Duden, R., Emr, S.D., Riezman, H., Cosson, P., 1994. Coatomer is essential for retrieval of dilysine-tagged proteins to the endoplasmic reticulum. Cell 79, 1199–1207. 10.1016/0092-8674(94)90011-6

Levtchenko, E.N., Wilmer, M.J.G., Janssen, A.J.M., Koenderink, J.B., Visch, H.J., Willems, P.H.G.M., De Graaf-Hess, A., Blom, H.J., Van Den Heuvel, L.P., Monnens, L.A., 2006. Decreased intracellular ATP content and intact mitochondrial energy generating capacity in human cystinotic fibroblasts. Pediatr. Res. 59, 287–292. 10.1203/01.pdr.0000196334.46940.54

Li, C., Dougherty, L., Coluccio, A.E., Meng, D., El-Sharkawy, I., Borejsza-Wysocka, E., Liang, D., Piñeros, M.A., Xu, K., Cheng, L., 2020. Apple ALMT9 Requires a Conserved C-Terminal Domain for Malate Transport Underlying Fruit Acidity. Plant Physiol. 182, 992–1006. 10.1104/pp.19.01300

Liu, B., Du, H., Rutkowski, R., Gartner, A., Wang, X., 2012. LAAT-1 is the lysosomal lysine/arginine transporter that maintains amino acid homeostasis. Science 337, 351–354. 10.1126/science.1220281

Löbel, M., Salphati, S.P., El Omari, K., Wagner, A., Tucker, S.J., Parker, J.L., Newstead, S., 2022. Structural basis for proton coupled cystine transport by cystinosin. Nature Communications 2022 13:1 13, 1–12. 10.1038/s41467-022-32589-2

Lobry, T., Miller, R., Nevo, N., Rocca, C.J., Zhang, J., Catz, S.D., Moore, F., Thomas, L., Pouly, D., Bailleux, A., Guerrera, I.C., Gubler, M.C., Cheung, W.W., Mak, R.H., Montier, T., Antignac, C., Cherqui, S., 2019. Interaction between galectin-3 and cystinosin uncovers a pathogenic role of inflammation in kidney involvement of cystinosis. Kidney Int. 96, 350–362. 10.1016/j.kint.2019.01.029

Lohman, J.R., Remington, S.J., 2008. Development of a family of redox-sensitive green fluorescent protein indicators for use in relatively oxidizing subcellular environments. Biochemistry 47, 8678–8688. 10.1021/BI800498G/SUPPL_FILE/BI800498G-FILE002.PDF

Lord, C., Ferro-Novick, S., Miller, E.A., 2013. The highly conserved COPII coat complex sorts cargo from the endoplasmic reticulum and targets it to the golgi. Cold Spring Harb. Perspect. Biol. 5. 10.1101/cshperspect.a013367

Luciani, A., Devuyst, O., 2024. The CTNS-MTORC1 axis couples lysosomal cystine to epithelial cell fate decisions and is a targetable pathway in cystinosis. Autophagy 20, 202–204. 10.1080/15548627.2023.2250165

Luciani, A., Festa, B.P., Chen, Z., Devuyst, O., 2018. Defective autophagy degradation and abnormal tight junction-associated signaling drive epithelial dysfunction in cystinosis. Autophagy 14, 1157–1159. 10.1080/15548627.2018.1446625

Luo, M., Jiang, Y.L., Ma, X.X., Tang, Y.J., He, Y.X., Yu, J., Zhang, R.G., Chen, Y., Zhou, C.Z., 2010. Structural and Biochemical Characterization of Yeast Monothiol Glutaredoxin Grx6. J. Mol. Biol. 398, 614–622. 10.1016/j.jmb.2010.03.029

Manna, P.T., Davis, L.J., Robinson, M.S., 2019. Fast and cloning-free CRISPR/Cas9-mediated genomic editing in mammalian cells. Traffic 20, 974–982. 10.1111/tra.12696

Mannucci, L., Pastore, A., Rizzo, C., Piemonte, F., Rizzoni, G., Emma, F., 2006. Impaired activity of the gamma-glutamyl cycle in nephropathic cystinosis fibroblasts. Pediatr. Res. 59, 332–335. 10.1203/01.pdr.0000196370.57200.da

Mesecke, N., Spang, A., Deponte, M., Herrmann, J.M., 2008. A novel group of glutaredoxins in the cis-Golgi critical for oxidative stress resistance. Mol. Biol. Cell 19, 2673–2680. 10.1091/MBC.E07-09-0896

Mirdita, M., Schütze, K., Moriwaki, Y., Heo, L., Ovchinnikov, S., Steinegger, M., 2022. ColabFold: making protein folding accessible to all. Nature Methods 2022 19:6 19, 679–682. 10.1038/s41592-022-01488-1

Napolitano, G., Johnson, J.L., He, J., Rocca, C.J., Monfregola, J., Pestonjamasp, K., Cherqui, S., Catz, S.D., 2015. Impairment of chaperone-mediated autophagy leads to selective lysosomal degradation defects in the lysosomal storage disease cystinosis. EMBO Mol. Med. 7, 158. 10.15252/EMMM.201404223

Newstead, S., Barr, F., 2020. Molecular basis for KDEL-mediated retrieval of escaped ER-resident proteins - SWEET talking the COPs. J. Cell Sci. 133. 10.1242/jcs.250100

Outten, C.E., Albetel, A.N., 2013. Iron sensing and regulation in Saccharomyces cerevisiae: Ironing out the mechanistic details. Curr. Opin. Microbiol. 16, 662–668. 10.1016/j.mib.2013.07.020

Pisoni, R.L., Thoene, J.G., Christensen, H.N., 1985. Detection and characterization of carrier-mediated cationic amino acid transport in lysosomes of normal and cystinotic human fibroblasts. Role in therapeutic cystine removal? Journal of Biological Chemistry 260, 4791–4798. 10.1016/s0021-9258(18)89141-4

Pollard, M.G., Travers, K.J., Weissman, J.S., 1998. Ero1p: A novel and ubiquitous protein with an essential role in oxidative protein folding in the endoplasmic reticulum. Mol. Cell 1, 171–182. 10.1016/S1097-2765(00)80018-0

Puigpinós, J., Casas, C., Herrero, E., 2015. Altered intracellular calcium homeostasis and endoplasmic reticulum redox state in Saccharomyces cerevisiae cells lacking Grx6 glutaredoxin. Mol. Biol. Cell 26, 104–116. 10.1091/MBC.E14-06-1137

Pujol-Carrion, N., Belli, G., Herrero, E., Nogues, A., de la Torre-Ruiz, M.A., 2006. Glutaredoxins Grx3 and Grx4 regulate nuclear localisation of Aft1 and the oxidative stress response in Saccharomyces cerevisiae. J. Cell Sci. 119, 4554–4564. 10.1242/jcs.03229

Rega, L.R., Polishchuk, E., Montefusco, S., Napolitano, G., Tozzi, G., Zhang, J., Bellomo, F., Taranta, A., Pastore, A., Polishchuk, R., Piemonte, F., Medina, D.L., Catz, S.D., Ballabio, A., Emma, F., 2016. Activation of the transcription factor EB rescues lysosomal abnormalities in cystinotic kidney cells. Kidney Int. 89, 862–873. 10.1016/j.kint.2015.12.045

Ritchie, M.E., Phipson, B., Wu, D., Hu, Y., Law, C.W., Shi, W., Smyth, G.K., 2015. limma powers differential expression analyses for RNA-sequencing and microarray studies. Nucleic Acids Res. 43, e47–e47. 10.1093/nar/gkv007

Ruivo, R., Bellenchi, G.C., Chen, X., Zifarelli, G., Sagneá, C., Debacker, C., Pusch, M., Supplisson, S., Gasnier, B., 2012. Mechanism of proton/substrate coupling in the heptahelical lysosomal transporter cystinosin. Proc. Natl. Acad. Sci. U. S. A. 109. 10.1073/pnas.1115581109

Sansanwal, P., Li, L., Hsieh, S.C., Sarwal, M.M., 2010. Insights into novel cellular injury mechanisms by gene expression profiling in nephropathic cystinosis. J. Inherit. Metab. Dis. 33, 775–786. 10.1007/S10545-010-9203-6

Sardana, R., Zhu, L., Emr, S.D., 2019. Rsp5 Ubiquitin ligase–mediated quality control system clears membrane proteins mistargeted to the vacuole membrane. Journal of Cell Biology. 10.1083/jcb.201806094

Sardana, R., Emr, S.D., 2021. Membrane protein quality control mechanisms in the endo-lysosome system. Trends in Cell Biology. 10.1016/j.tcb.2020.11.011

Sardana, R., Highland, C.M., Straight, B.E., Chavez, C.F., Fromme, J.C., Emr, S.D., 2021. Golgi membrane protein Erd1 Is essential for recycling a subset of Golgi glycosyltransferases. Elife 10. 10.7554/ELIFE.70774

Saudek, V., 2017. Potential dual function of PQ-loop proteins such as cystinosin. Journal of Biological Chemistry 292, 15133. 10.1074/jbc.L117.795278

Saudek, V., 2012. Cystinosin, MPDU1, SWEETs and KDELR Belong to a Well-Defined Protein Family with Putative Function of Cargo Receptors Involved in Vesicle Trafficking. PLoS One 7, e30876. 10.1371/JOURNAL.PONE.0030876

Schenk, B., Imbach, T., Frank, C.G., Grubenmann, C.E., Raymond, G. V., Hurvitz, H., Raas-Rotschild, A., Luder, A.S., Jaeken, J., Berger, E.G., Matthijs, G., Hennet, T., Aebi, M., 2001. MPDU1 mutations underlie a novel human congenital disorder of glycosylation, designated type If. Journal of Clinical Investigation 108, 1687. 10.1172/jci13419

Schmidt, O., Weyer, Y., Baumann, V., Widerin, M.A., Eising, S., Angelova, M., Schleiffer, A., Kremser, L., Lindner, H., Peter, M., Fröhlich, F., Teis, D., 2019. Endosome and Golgi-associated degradation (EGAD) of membrane proteins regulates sphingolipid metabolism. EMBO J. 38, e101433. 10.15252/EMBJ.2018101433

Schmitz, K.R., Liu, J., Li, S., Setty, T.G., Wood, C.S., Burd, C.G., Ferguson, K.M., 2008. Golgi localization of glycosyltransferases requires a Vps74p oligomer. Dev. Cell 14, 523–534. 10.1016/J.DEVCEL.2008.02.016

Schneider, J.A., Bradley, K., Seegmiller, J.E., 1967. Increased Cystine in Leukocytes from Individuals Homozygous and Heterozygous for Cystinosis. Science (1979). 157, 1321–1322. 10.1126/SCIENCE.157.3794.1321

Sevier, C.S., Kaiser, C.A., 2008. Ero1 and redox homeostasis in the endoplasmic reticulum. Biochim. Biophys. Acta Mol. Cell Res. 1783, 549–556. 10.1016/j.bbamcr.2007.12.011

Simpkins, J.A., Rickel, K.E., Madeo, M., Ahlers, B.A., Carlisle, G.B., Nelson, H.J., Cardillo, A.L., Weber, E.A., Vitiello, P.F., Pearce, D.A., Vitiello, S.P., 2016. Disruption of a cystine transporter downregulates expression of genes involved in sulfur regulation and cellular respiration. Biol. Open 5, 689–697. 10.1242/BIO.017517

Stokes, A.J., Wakano, C., Del Carmen, K.A., Koblan-Huberson, M., Turner, H., 2005. Formation of a physiological complex between TRPV2 and RGA protein promotes cell surface expression of TRPV2. J. Cell. Biochem. 94, 669–683. 10.1002/jcb.20331

Sumayao, R., McEvoy, B., Newsholme, P., McMorrow, T., 2016. Lysosomal cystine accumulation promotes mitochondrial depolarization and induction of redox-sensitive genes in human kidney proximal tubular cells. J. Physiol. 594, 3353–3370. 10.1113/JP271858

Talaia, G., Amick, J., Ferguson, S.M., 2021. Receptor-like role for PQLC2 amino acid transporter in the lysosomal sensing of cationic amino acids. Proc. Natl. Acad. Sci. U. S. A. 118. 10.1073/pnas.2014941118

Taranta, A., Bellomo, F., Petrini, S., Polishchuk, E., De Leo, E., Rega, L.R., Pastore, A., Polishchuk, R., De Matteis, M.A., Emma, F., 2016. Cystinosin-LKG rescues cystine accumulation and decreases apoptosis rate in cystinotic proximal tubular epithelial cells. Pediatr. Res. 81, 113–119. 10.1038/PR.2016.184

Taranta, A., Petrini, S., Citti, A., Boldrini, R., Corallini, S., Bellomo, F., Levtchenko, E., Emma, F., 2012. Distribution of cystinosin-LKG in human tissues. Histochem. Cell Biol. 138, 351–363. 10.1007/S00418-012-0958-8

Taranta, A., Petrini, S., Palma, A., Mannucci, L., Wilmer, M.J., De Luca, V., Diomedi-Camassei, F., Corallini, S., Bellomo, F., Van Den Heuvel, L.P., Levtchenko, E.N., Emma, F., 2008. Identification and subcellular localization of a new cystinosin isoform. Am. J. Physiol. Renal Physiol. 294. 10.1152/AJPRENAL.00413.2007

Taylor, R.J., Zubkov, N., Ciazynska, K.A., Kaufman, J.G.G., Tagiltsev, G., Owen, D.J., Briggs, J.A.G., Munro, S., 2025. The mechanistic basis of cargo selection during Golgi maturation. Sci. Adv. 11, 1–11. 10.1126/sciadv.aea0016

Thirumalaikumar, V.P., Wagner, M., Balazadeh, S., Skirycz, A., 2021. Autophagy is responsible for the accumulation of proteogenic dipeptides in response to heat stress in Arabidopsis thaliana. FEBS Journal 288, 281–292. 10.1111/febs.15336

Town, M., Jean, G., Cherqui, S., Attard, M., Forestier, L., Whitmore, S.A., Gallen, D.F., Gribouval, O., Broyer, M., Bates, G.P., Hoff, W.V. t., Antignac, C., 1998. A novel gene encoding an integral membrane protein is mutated in nephropathic cystinosis. Nature Genetics 1998 18:4 18, 319–324. 10.1038/ng0498-319

Tu, L., Tai, W.C.S., Chen, L., Banfield, D.K., 2008. Signal-mediated dynamic retention of glycosyltransferases in the Golgi. Science 321, 404–407. 10.1126/SCIENCE.1159411

Wang, J., Sevier, C.S., 2016. Formation and Reversibility of BiP Protein Cysteine Oxidation Facilitate Cell Survival during and post Oxidative Stress. J. Biol. Chem. 291, 7541. 10.1074/JBC.M115.694810

Wang, Y., Menon, A.K., Maki, Y., Liu, Y.S., Iwasaki, Y., Fujita, M., Guerrero, P.A., Silva, D.V. n., Seeberger, P.H., Murakami, Y., Kinoshita, T., 2022. Genome-wide CRISPR screen reveals CLPTM1L as a lipid scramblase required for efficient glycosylphosphatidylinositol biosynthesis. Proceedings of the National Academy of Sciences 119, e2115083119. 10.1073/pnas.2115083119

Wei, H., Kim, S.J., Zhang, Z., Tsai, P.C., Wisniewski, K.R., Mukherjee, A.B., 2008. ER and oxidative stresses are common mediators of apoptosis in both neurodegenerative and non-neurodegenerative lysosomal storage disorders and are alleviated by chemical chaperones. Hum. Mol. Genet. 17, 469–477. 10.1093/hmg/ddm324

Weyer, Y., Schwabl, S.I., Tang, X., Purwar, A., Siegmann, K., Ruepp, A., Dunzendorfer-Matt, T., Widerin, M.A., Niedrist, V., Mutsters, N.J.M., Tettamanti, M.G., Weys, S., Sarg, B., Kremser, L., Liedl, K.R., Schmidt, O., Teis, D., 2024. The Dsc ubiquitin ligase complex identifies transmembrane degrons to degrade orphaned proteins at the Golgi. Nat. Commun. 15. 10.1038/s41467-024-53676-6

Wilmer, M.J., Kluijtmans, L.A.J., van der Velden, T.J., Willems, P.H., Scheffer, P.G., Masereeuw, R., Monnens, L.A., van den Heuvel, L.P., Levtchenko, E.N., 2011. Cysteamine restores glutathione redox status in cultured cystinotic proximal tubular epithelial cells. Biochim. Biophys. Acta Mol. Basis Dis. 1812, 643–651. 10.1016/j.bbadis.2011.02.010

Wilmer, M.J.G., De Graaf-Hess, A., Blom, H.J., Dijkman, H.B.P.M., Monnens, L.A., Van Den Heuvel, L.P., Levtchenko, E.N., 2005. Elevated oxidized glutathione in cystinotic proximal tubular epithelial cells. Biochem. Biophys. Res. Commun. 337, 610–614. 10.1016/j.bbrc.2005.09.094

Zhang, J., Johnson, J.L., He, J., Napolitano, G., Ramadass, M., Rocca, C., Kiosses, W.B., Bucci, C., Xin, Q., Gavathiotis, E., Cuervo, A.M., Cherqui, S., Catz, S.D., 2017. Cystinosin, the small GTPase Rab11, and the Rab7 effector RILP regulate intracellular trafficking of the chaperone-mediated autophagy receptor LAMP2A. Journal of Biological Chemistry 292, 10328–10346. 10.1074/jbc.M116.764076

Zhou, Y., Liu, Y., Gupta, S., Paramo, M.I., Hou, Y., Mao, C., Luo, Y., Judd, J., Wierbowski, S., Bertolotti, M., Nerkar, M., Jehi, L., Drayman, N., Nicolaescu, V., Gula, H., Tay, S., Randall, G., Wang, P., Lis, J.T., Feschotte, C., Erzurum, S.C., Cheng, F., Yu, H., 2022. A comprehensive SARS-CoV-2–human protein–protein interactome reveals COVID-19 pathobiology and potential host therapeutic targets. Nature Biotechnology 2022 41:1 41, 128–139. 10.1038/s41587-022-01474-0

